# Lost in communication: How Müller glia cells fail to maintain retinal integrity in USH1C retinal organoids

**DOI:** 10.1101/2025.06.03.657614

**Authors:** Nicole Wenck, Mark Zorin, Qiang Wang, Selina Kröll, Katharina Hay, Christof Rickert, Maria Méndez-Lago, Sivarajan Karunanithi, Dimitra Athanasiou, Kalliopi Ziaka, Pietro De Angeli, Katarina Stingl, Susanne Kohl, Michael E. Cheetham, Thomas Mittmann, Kerstin Nagel-Wolfrum

**Affiliations:** Institute of Developmental Biology and Neurobiology, Johannes Gutenberg-University Mainz, Mainz, Germany; Institute of Molecular Physiology, Johannes Gutenberg-University Mainz, Mainz, Germany; Institute of Physiology, University Medical Centre of Johannes Gutenberg-University Mainz, Mainz, Germany; Light Microscopy Core Facility (LMCF) of Faculty of Biology, Johannes Gutenberg-University Mainz, Mainz, Germany; Institute of Molecular Biology GmbH (IMB), Mainz, Germany; UCL Institute of Ophthalmology, University College London, London, UK; Institute for Ophthalmic Research, Centre for Ophthalmology, University of Tuebingen, Tuebingen, Germany; University Eye Hospital, Centre for Ophthalmology, University of Tuebingen, Tuebingen, Germany; Institute of Quantitative and Computational Biosciences (IQCB), Johannes Gutenberg-University Mainz, Mainz, Germany

**Keywords:** Inherited retinal dystrophy, *USH1C*, human retinal organoids, disease pathology, intercellular communication, outer limiting membrane

## Abstract

Usher syndrome type 1, caused by pathogenic variants in the *USH1C* gene, leads to congenital deafness and progressive retinal degeneration resulting in vision loss. While auditory deficits can be compensated by cochlea implants and hearing aids, no treatment exists to prevent retinal degeneration. Here, we generated retinal organoids from induced pluripotent stem cells of two USH1C patients to elucidate the cellular and molecular mechanisms driving ocular pathogenesis. Single-cell RNA sequencing of both healthy and USH1C retinal organoids identified differential expression of genes related to phototransduction in photoreceptors, as well as alterations in cell adhesion and canonical Wnt signaling in Müller glia cells. Analysis of intercellular communication revealed an overall reduced signaling efficiency, particularly affecting Müller glia-mediated retinal adhesion processes. Morphological characterization of organoids confirmed transcriptome changes by showing degeneration of the outer limiting membrane and loss of adherens junction architecture. Moreover, photoreceptors revealed increased susceptibility to morphological and functional changes related to phototransduction. These results demonstrate that disruption of Müller glia signaling contributes to a loss of retinal integrity, providing novel insights into USH1C pathogenesis and offering targets for therapeutic interventions.

## Introduction

Usher syndrome is an autosomal recessive disorder characterized by sensorineural hearing loss, *retinitis pigmentosa* (RP) and vestibular dysfunction. With 1:5,000 to 1:10,000 affected individuals, it is the most common cause of inherited deaf-blindness [1]. Depending on severity and progression, the heterogenous disorder is categorized into three clinical subtypes (USH1, USH2, USH3), with a fourth subtype (USH4) under discussion [2]. Of those, USH1 is the most severe form, as patients are profoundly deaf from birth, have early childhood balance issues and prepubertal vision impairments. Vision loss primarily begins with a progressive degeneration of rod photoreceptors in the peripheral retina leading to night blindness and tunnel vision. Secondarily, degeneration of cone photoreceptors impairs central vision and eventually results in blindness [3].

*USH1C* is one of six genes characterized and linked to USH1. Located on chromosome 11p15.1, the gene comprises 28 exons and encodes the protein harmonin [4]. In humans, extensive alternative splicing of *USH1C* results in at least 11 different isoforms that fulfill diverse functions in various tissues, including the inner ear and retina [2, 4, 5]. While murine models demonstrated harmonin’s critical role in mechano-transduction within cochlear hair cells of the inner ear [6–8], its function in the retina remains poorly understood. A major reason is that Ush1c mice are not suitable for studying retinal pathology, as they have a different retinal structure to humans and often lack signs of visual degeneration [1, 9, 10]. Studies in alternative models demonstrated harmonin localization in two retinal cell types: photoreceptor and Müller glia (MG) cells [4, 11, 12]. While photoreceptors are essential for detecting light and visual signal processing, MG cells play a crucial role in the structural and metabolic support of retinal neurons and in maintaining retinal homeostasis [13, 14]. In both cell types, harmonin is thought to function as a scaffold protein, organizing and interacting with other proteins, which in turn fulfil different functions. Its binding partners include several USH proteins, such as cadherin 23 (*USH1D*), protocadherin 15 (*USH1F*) or usherin (*USH2A*) that interact with harmonin’s PDZ domains [7, 15–19]. Additionally, non-USH proteins bind to harmonin’s PDZ domains [4, 20–22]. Recent studies focused on the interaction between harmonin and the multifunctional protein β-catenin (*CTNNB1*), revealing two functions: first, their interaction in the adherens junction between photoreceptor and MG cells at the outer limiting membrane (OLM) is important for maintaining retinal integrity [4]. And second, their cytoplasmatic interaction negatively regulates canonical Wnt signaling, which is likely essential for retinal development and homeostasis [23]. Nevertheless, a more comprehensive knowledge of harmonin’s role in the retina is needed for developing targeted therapies for USH1C patients, as there is currently no treatment that slows or prevents retinal degeneration [2].

Recently, the generation of human induced pluripotent stem cell (hiPSC)-derived retinal organoids (ROs) has led to advances in the understanding of retinal development, retinal degeneration and the testing of potential therapies [24]. By encompassing all major neuroretinal cell types, these three-dimensional *in vitro* models recapitulate the cellular composition and stratified architecture of the native retina [25, 26]. Transcriptomic and proteomic analyses revealed that mature ROs exhibit expression profiles similar to those of the fetal human retina, thereby recapitulating key aspects of *in vivo* retinogenesis [25, 27–29]. Moreover, electrophysiological studies have demonstrated the functional capacity of RO photoreceptors to process and respond to light stimuli, which further highlights their physiological relevance [30–32]. With being derived from patient-based hiPSCs, ROs also offer the opportunity to investigate retinal pathology within a patient-specific genetic background. To date, various studies have been performed examining the molecular mechanisms of retinal degeneration and testing different therapeutic interventions in ROs [33–40].

In this study, we present a comprehensive analysis of the first USH1C RO model derived from hiPSCs of two siblings. Using single-cell RNA sequencing (scRNA-seq) we showed severe dysregulation of MG-mediated intercellular communication and differential gene expression related to retinal adhesion. Furthermore, we demonstrate OLM thinning and disrupted adherens junctions by immunofluorescence (IF)-analysis, which can be accompanied by transcriptional, morphological and functional impairments in photoreceptor cells. Overall, our RO model provides valuable insights into USH1C-associated retinal pathology and paves the way for future therapeutic approaches.

## Materials and methods

### Human dermal fibroblast cell culture

Dermal fibroblasts were expanded from skin biopsies of human subjects in DMEM + GlutaMAX (Thermo Fisher Scientific), 10% fetal bovine serum (FBS) (Cytiva) at 37°C, 5% CO_2_. Genetic characterization of fibroblasts from two USH1C patients was done by confirming the biallelic *USH1C* pathogenic variants c.91C>T; p.(R31*) and c.238dupC; p.(R80Pfs*69) [4].

### Human induced pluripotent stem cell generation and culture

Episomal reprogramming of fibroblasts was performed according to an established protocol [41]. Briefly, fibroblasts were dissociated using Trypsin (Thermo Fisher Scientific) and pelleted at 200 × g for 5 min. 1x10^6^ cells were resuspended in 100 µl P2 Primary Cell Nucleofector Solution (Lonza) mixed with 1 µg of each plasmid pCXLE-hSK, pCXLE-hUL, pCXLE-hOCT3/4 and pSIMPLE-miR302/367 (all: Addgene). After transferring fibroblasts to a Nucleocuvette vessel, cells were electroporated by 4D-Nucleofector program DT-130 (Lonza) and seeded on 0.2% gelatin-coated (Carl Roth) petri dishes. For the next 5 days, the media was replaced daily with DMEM + GlutaMAX, 10% FBS, 0.5 mM sodium butyrate (SB) (Sigma-Aldrich). At day 7, cells were dissociated using Trypsin and 200,000 cells per well were seeded onto a Geltrex-coated (Thermo Fisher Scientific) 6-well plate. Following, the media was switched to mTeSR Plus (StemCell Technologies), 0.5 mM SB from day 8 to 12. From day 13 on, cells were cultured in mTeSR Plus until colonies formed (approximately day 25), which were mechanically dissected and further expanded. Selected hiPSC clones were maintained on Geltrex coating and in mTeSR plus at 37°C, 5% CO_2_. At 70-80% confluency, hiPSCs were washed with 1 ml Dulbecco’s Phosphate Buffered Saline (DPBS) (Thermo Fisher Scientific) and detached using Cell Dissociation Buffer (Thermo Fisher Scientific) for 5 min at room temperature (RT). After aspirating Cell Dissociation Buffer, cells were harvested in 1 ml mTeSR plus, cell aggregates were diminished by gentle pipetting and hiPSCs were seeded on Geltrex-coated plates.

### Genomic stability of hiPSCs

Genomic stability of hiPSCs was assessed by detection of recurrent genetic abnormalities using the iCS-digital^TM^ PSC test, provided as a service by Stem Genomics (https://www.stemgenomics.com/), as described previously [42].

### Sanger sequencing of hiPSCs

Genomic DNA was isolated from hiPSCs using the peqGOLD Tissue DNA Mini Kit (Avantor). Exon 2 and 3 of the *USH1C* gene were partially amplified via PCR with specific primers (Table S1). The PCR product was purified using a PCR & DNA Cleanup Kit (New England Biolabs) and sequenced in a 3730 DNA Analyzer (StarSEQ, Mainz, Germany).

### Embryoid body assay

Embryoid body (EB)-assay was conducted as previously reported with slight modifications [43]. Dissociation of hiPSCs was performed using Cell Dissociation Buffer as described above. Following, single hiPSCs were seeded on ultra-low attachment dishes containing mTeSR plus, 10 µM Y27632 (StemCell Technologies). On day 3, the media was switched to DMEM/F-12 + GlutaMAX, 20% Knockout Serum Replacement, 1% Non-Essential Amino Acids (NEAA), 1% GlutaMAX, 55 mM β-mercaptoethanol (all: Thermo Fisher Scientific). After 7 days, EBs were seeded on Geltrex-coated coverslips and cultured in the same medium. Samples were collected for IF-analysis on day 17.

### Immunofluorescence analysis of hiPSCs

IF-analysis of hiPSCs was performed as previously reported [44]. hiPSCs were cultured on Geltrex-coated coverslips. Cells were washed with DPBS once and fixed with 4% paraformaldehyde (PFA) (Merck) for 10 min at RT. PFA was removed, cells were washed with DPBS twice and incubated in DPBS, 1% BSA (Carl Roth), 0.3% Triton X-100 (Sigma-Aldrich) for 15 min at RT. After permeabilization and blocking, primary antibodies (Table S2) were diluted in DPBS, 1% BSA, 0.3% Triton X-100 and applied overnight at 4 °C. Following, cells were washed with DPBS twice and incubated with secondary antibodies (Table S3) diluted in 1% BSA, 0.3% Triton X-100 for 2 h at RT. Embedding was performed using Mowiol (Roth) after washing cells with DPBS twice. Images were taken using a Leica DM 6000 fluorescence microscope.

### Retinal organoid differentiation

ROs were generated as previously described with slight modifications [45]. Differentiation started at day 0, when hiPSCs reached 70-80% confluency and were transferred to Essential 6 medium (E6) (Thermo Fisher Scientific) for 2 days. On day 3, the media was changed to E6, 1x N2 supplement (Thermo Fisher Scientific) and replaced three times a week. Emerging neuronal vesicles were manually cut using disposable scalpels (Fisher Scientific) and transferred to ultralow attachment plates (Thermo Fisher Scientific) at day 28. Medium was switched to DMEM/F12 + GlutaMAX, 2% B27 supplement, 1% GlutaMAX, 1% MEM non-essential amino acids (NEAA), 0.1% Penicillin/Streptomycin (Pen/Strep) (all: Thermo Fisher Scientific), 15 ng/ml fibroblast growth factor 2 (FGF2) (Miltenyi Biotech). From day 35, medium was replaced two times a week and switched to DMEM/F12 + GlutaMAX, 10% FBS, 2% B27 supplement, 1% GlutaMAX, 1% NEAA, 0.1% Pen/Strep, 100 mM Taurine (Sigma-Aldrich). From day 65 on, the basis medium consisting of DMEM/F12 + GlutaMAX, 10% FBS, 2% B27 supplement without vitamin A (Thermo Fisher Scientific), 1% GlutaMAX, 1% NEAA, 0.1% Pen/Strep was used. Until day 85, 100 mM Taurine and 1 mM Retinoic Acid (Sigma-Aldrich) were supplemented. Additional 1x N2 supplement was added from day 85 until day 120. From day 120 on, 1x N2 and 100 mM Taurine was added. RO growth was regularly monitored during the differentiation using an EVOS XL Core microscope (Thermo Fisher Scientific).

### Single cell dissociation of ROs

For multiplexed scRNA-seq four healthy-1 and three USH1C-1 ROs were processed. Single cell dissociation was performed as previously reported with slight modifications [25]. Briefly, ROs were washed twice with 1 ml pre-warmed Hank’s Balanced Salt Solution (HBSS) (Thermo Fisher Scientific). For dissociation, ROs were incubated with 300 µl activated papain solution containing 8 U papain (CellSystems) and 48 µl activator consisting of 1.1 µM EDTA (Sigma-Aldrich), 5.5 mM L-cysteine (Fisher Scientific), 0.07 mM 2-mercaptoethanol (Thermo Fisher Scientific) and H_2_O for 40 min at 37°C, 5% CO_2_. Tubes were gently inverted every 10 min and dissociation was stopped by placing the tubes on ice and adding 300 µl stop solution per RO containing DMEM/F12 + GlutaMAX, 10% FBS and 20 U/ml DNase (Sigma-Aldrich). After centrifuging at 200 × g and 4 °C for 30 s, ROs were washed with 1 ml DMEM/F12 + GlutaMAX, 10% FBS. Mechanical dissociation was performed by gently pipetting 20 times in 300 µl DMEM/F12 + GlutaMAX, 10% FBS until no clumps were visible anymore.

### Single cell RNA-sequencing library preparation and sequencing

Fixation of single cells was performed by using the Chromium Next GEM Single Cell Fixed RNA Sample Preparation Kit (10X Genomics) according to the manufacturer’s instructions. For probe hybridization and barcoding, we followed the guidelines (CG000527) of the Chromium Fixed RNA Profiling Reagent Kits, Human Transcriptome (10X Genomics). Between 284,000 and 824,000 cells per sample were used for hybridization to probe sets BC001-BC007. On average, 15,300 cells per sample were pooled for subsequent gel bead in emulsion (GEM) generation and gene expression library preparation. Library sequencing was performed in the NextSeq 2000 sequencer (Illumina), on a P3 100 flow cell, with 28 cycles for Read 1 and 90 cycles for Read 2, at the Genomics Core Facility, Institute of Molecular Biology, Mainz.

### Single cell RNA-sequencing data processing and quality control

Sequence quality of FastQ files was assessed via FastQC (v0.11.9) [46]. The samples were demultiplexed and analyzed using the 10X Genomics Cell Ranger software (v7.0.1; pipeline: multi) [47]. Reads were then mapped against reference genome GRCh38 (ensembl98, refdata-gex-GRCh38-2020-A) and counts were quantified using the probe set (Chromium_Human_Transcriptome_Probe_Set_v1.0_GRCh38-2020-A) provided by 10XGenomics with default parameters. As the samples were multiplexed with other project data, sample-specific fastq files related to this manuscript were generated from the aligned BAM files using the cellranger bamtofastq pipeline (https://github.com/10XGenomics/bamtofastq). Downstream analysis was performed in a Conda environment (v24.7.1) using R (v4.3.3).

Quality control of cells and downstream analyses were conducted using the Seurat package (v5.1.0) [48]. Cells with mitochondrial gene expression levels three standard deviations above or below the median for each sample were removed. Cells whose numbers of detected transcripts were three standard deviations above or below the median per sample were excluded. Based on these quality control thresholds, 4.2% of cells were removed. Following, normalization was done using the SCTransform function (v2) of Seurat package (v0.4.1) [49]. Samples were integrated using IntegrateData and FindIntegrationAnchors function with CCA as a reduction method.

### Clustering and cell type annotation

On a reduced 30-dimensional space obtained via the RunPCA function (npcs = 50), a shared nearest neighbour graph was constructed using the FindNeighbors function (k.param = 20; ndims = 1:30). Using the above graph, cells were clustered using the original Louvain algorithm as implemented in the FindClusters function with a resolution of 1.6. Subsequently, clusters were sub-clustered via the FindSubCluster function with a resolution of 0.2. After clustering, the sequencing depth was fixed for all cells as SCT assay data using PrepSCTFindMarkers function. SCT assay data were used for all downstream analyses. For cluster annotation, cluster-specific differentially expressed genes were identified via FindAllMarkers function using MAST algorithm (v1.28.0) [50]. Additionally, literature-derived marker genes were used to annotate the clusters [51–54]. For all steps, apart from the parameters mentioned, default settings were used.

### Composition analysis

To compare the distribution of cell types between healthy-1 and USH1C-1 ROs, a proportional analysis was performed using sccomp package (v1.99.18) [55]. The linear model was fitted with the sccomp_estimate function, followed by the sccomp_test function. Proportional changes were calculated with the sccomp_proportional_fold_change function, using healthy-1 ROs as the reference group.

### Pseudotime analysis

Pseudotime analysis was performed using the Monocle 3 package (v1.3.7) to characterize the trajectory of gene expression changes [56]. Amacrine, horizontal and retinal pigment epithelial (RPE) cells were excluded from analysis due to their cluster separation reflecting their early terminal differentiation state. Other clusters were connected to form a single pseudotime trajectory. Early retinal precursor cells (RPCs) were manually set as the starting point of differentiation.

### CellChat analysis

Intercellular communication was analyzed using CellChat package (v2.1.2) [57]. CellChat objects were created separately for healthy-1 and USH1C-1 ROs from SCT data using the createCellChat function. To identify cell-cell communication profiles, we utilized the complete CellChatDB.human database of ligand-receptor interactions. CellChat pipeline was used with default parameters, unless stated otherwise. Overexpressed ligand-receptor interactions were identified via identifyOverExpressedGenes and identifyOverExpressedInteractions functions. After that, cell-cell communication profiles were computed using computeCommunProb function with parameter population.size = FALSE. Filtration of cell-cell communication as well as computation of communication probability and aggregated communication network were done using filterCommunication, computeCommunProbPathway and aggregateNet functions. Finally, CellChat objects of healthy-1 and USH1C-1 ROs were merged via mergeCellChat function.

### Differential gene expression analysis

Condition-specific differential expressed genes (DEGs) (healthy-1 vs. USH1C-1) were identified using FindMarkers funtion using SCT values and MAST algorithm with the parameters min.pct = 0.1 and pseudocount.use = 0.1. Non-significant DEGs were filtered out using an adjusted p-value (p.adjust) of 0.05. Enrichment analysis of DEGs was performed using Cytoscape (v3.9.1), ClueGO (v2.5.10) and CluePedia (v1.5.10). DEGs were enriched for the Gene Ontology (GO) categories Biological Process (EBI-UniProt-GOA-ACAP-ARAP-20.03.2024), Cellular Component (EBI-UniProt-GOA-ACAP-ARAP-20.03.2024), and for Kyoto Encyclopedia of Genes and Genomes (KEGG)-pathways (KEGG-20.03.2024). Significance testing for enrichment analysis was done using a two-sided hypergeometric test and adjusted for multiple hypothesis testing by Bonferroni correction. GO-/KEGG-terms with a p.adjust less than 0.05 were considered significant. Within each GO-/KEGG category, the five most significant GO-terms and the three most significant KEGG-terms were displayed. The ratio of Associated DEGs was defined as follows:

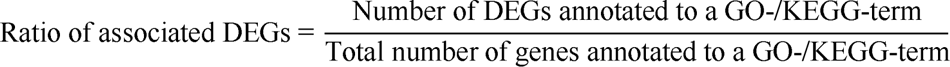

### Immunofluorescence analysis of ROs

IF-staining for ROs was conducted by using an established protocol [45]. Briefly, ROs were washed with DPBS once and fixed with 4% PFA for 20 min at 4 °C. After washing three times with DPBS, samples were incubated in 30% sucrose at 4 °C overnight. Following, ROs were embedded in Tissue-Tek O.C.T. (Fisher Scientific) and snap-frozen on dry ice. 10 µm thick cryosections were performed and mounted onto SuperFrost Plus Slides (Fisher Scientific). For immunostaining, cryosections were washed with DPBS once and incubated in DPBS, 10% donkey serum (Abcam), 5% BSA, 0.1% Triton X-100 for 1 h at RT. Primary antibodies (Table S2) were diluted in DPBS, 2% donkey serum, 1% BSA, 0.1% Triton X-100 and applied on cryosections overnight at 4 °C. After washing with DPBS three times, secondary antibodies (Table S3) were diluted in DPBS, 2% donkey serum, 1% BSA, 0.1% Triton X-100 and incubated on the sections for 2 h at RT. Embedding was performed using Mowiol after washing sections with DPBS three times. Imaging was performed using a ZEISS LSM 900 confocal microscope. For super-resolution imaging, Airyscan 2 was applied.

### Quantitative IF-analysis

To quantify the RHO-, GNAT1-, GUCA1A-, SAG- and ARR3-positive IS/OS area, maximum intensity projection was applied to all images. For measurements, only phototransduction protein-positive areas apical to the ONL were included. For this, basal parts were manually removed in Fiji. Images were converted into 8-bit format and thresholded using the default threshold mode. Following, the positive pixel area was measured and normalized to the RO circumference.

The total amount of DAPI-positive and SOX9-positive cells was assessed using a Python script (v3.9.21) in Jupyter notebook (v4.3.4) (see Data and code availability section). Mislocalized SOX9-positive MG cells were manually counted and defined as such when located in the ONL as indicated by NRL staining.

For quantification of F-actin and N-cadherin intensities at the OLM, all sections were imaged with the same laser intensity and exposure time. Maximum intensity projection was applied to the acquired images. Measurements were performed using a Fiji script (see Data and code availability section) on randomly selected regions of interest (ROIs) encompassing the OLM-specific staining apical to the ONL [58]. To ensure comparability, ROIs which included OLM-related pores (no staining) were excluded from analysis. For each image, the mean intensity was calculated from 10 measured ROIs and normalized to the mean intensity of three randomly selected background ROIs.

### Electrophysiology

Whole-cell patch-clamp recordings from single cone photoreceptor cells in ROs were performed according to an established protocol [31]. Shortly, ROs were placed in a submerged recording chamber, mounted on an upright Olympus BX51WI microscope and constantly perfused with oxygenated Ames’ solution at 37°C. Ames’ solution contained (in mM): 120 NaCl, 22.6 NaHCO_3_, 6 D-glucose, 3.1 KCl, 1.2 MgSO_4_, 1.1 CaCl_2_, 0.5 K_2_HPO_4_ and 0.5 L-glutamine. Borosilicate glass recording pipettes (Science Products) were pulled with a tip resistance of 3-6 MΩ using a DMZ Universal Electrode Puller (DMZ-Instruments) and filled with a pipette solution containing (in mM): 110 KCL, 13 NaCl, 2 MgCl_2_, 1 CaCl_2_, 10 EGTA, 10 HEPES, 5 ATP-Na, 0.5 GTP-Na, pH adjusted to 7.2 with KOH. The electrophysiological recordings were carried out using an Axopatch-200B amplifier connected to a Digidata 1440A and controlled by pClamp (v11.1) (all: Molecular Devices). Cone OS were identified based on their round structure and apical localization. Only smooth and bright cones were chosen for patch-clamp recordings. Access resistance was evaluated before and after each patch-clamp recording. Only cones with a relatively small access resistance change of <15% during the entire recording period were included for data analysis.

### Statistical analysis

Statistical analysis was conducted using GraphPad Prism (v10.4.0). For comparison of two groups, an unpaired two-tailed *t*-test or Wilcoxon rank-sum test was used, depending on the distribution and variance of data. For comparison among multiple groups, one-way or two-way ANOVA with Tukey’s post-hoc test was applied as appropriate. In case of unequal variances, the Kruskal-Wallis-test was conducted. Statistical analyses were performed on data from n ≥ 3 ROs per cell line. Data are presented as mean ± SD. P-values < 0.05 were considered significant.

## Results

### Characterization of USH1C-1 retinal organoids revealed altered outer limiting membrane development

Human dermal skin fibroblasts were collected from a healthy donor (healthy-1) and patient (USH1C-1), who harbors a nonsense variant c.91C>T; p.(R31*) and a 1 bp duplication c.238dupC; p.(R80Pfs*69) in the *USH1C* gene (Fig. S1A). The first pathogenic variant leads to an in-frame premature termination codon (PTC) in exon 2, while the second variant causes a frameshift in exon 3 and a PTC in exon 5 (Fig. S1B). We performed episomal reprogramming of healthy-1 and USH1C-1 fibroblasts into hiPSC and confirmed their respective *USH1C* variants via Sanger sequencing [41, 59, 60] (Fig. S1C; Fig. S2A, S2B). Copy number variation (CNV) analysis demonstrated genetic stability in both hiPSC lines (Fig. S2C). Pluripotency was validated through an embryoid body (EB) assay, which showed expression of germ layer markers Nestin (ectoderm), Brachyury (mesoderm), and AFP (endoderm) (Fig. S2D). The undifferentiated state of hiPSCs was further confirmed by the expression of stem cell transcription factors NANOG, OCT4, and SOX2 (Fig. S2E).

Next, we differentiated healthy-1 and USH1C-1 hiPSCs into ROs following an established protocol [45]. During the differentiation process, we transferred neuronal vesicles to three-dimensional culture and performed several media changes until ROs developed, which were considered mature at week 35 (w35) (Fig. 1A) [25]. To determine morphological differences during RO development, we monitored the final maturation stages by taking phase contrast images of healthy-1 and USH1C-1 ROs at four different time points (w20, w25, w30, w35) (Fig. 1B). Quantification of the brush border length revealed accelerated growth of photoreceptor inner and outer segments (IS/OS) in USH1C-1 ROs from w20 to w30. Interestingly, mature healthy-1 and USH1C-1 ROs showed no differences in IS/OS length at w35. As harmonin localizes in the adherens junction between photoreceptor and MG cells at the OLM, we additionally measured the OLM thickness over time (Fig. 1B). Both RO lines exhibited steady OLM growth from w20 to w30. However, from w30 to w35, the OLM width in USH1C-1 ROs decreased compared to healthy-1 ROs.

**Fig. 1.**
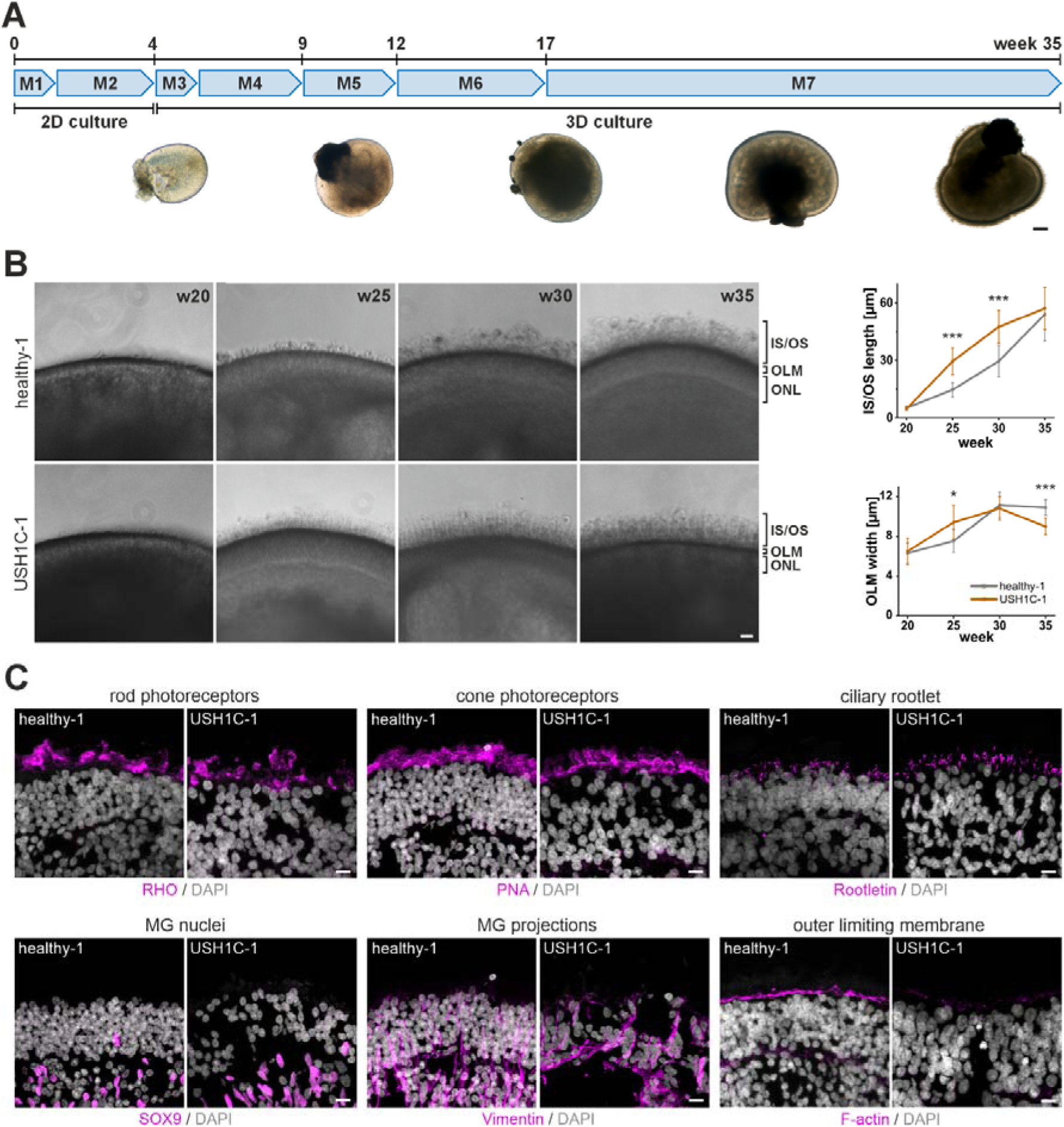
Morphological characterization of healthy-1 and USH1C-1 retinal organoids. **A)** Differentiation of hiPSCs into mature retinal organoids (ROs) takes 35 weeks and several media changes (M1-M7). Scale bar: 200 µm. **B)** Representative images showing the final 15 weeks of maturation for healthy-1 and USH1C-1 ROs. Scale bar: 25 µm. Quantification of the inner/outer segment (IS/OS) length and outer limiting membrane (OLM) width at week (w) 20, 25, 30 and 35. Independent samples *t*-test for n = 6-10 ROs per cell line/time point, 12 measurements per RO. **C)** IF-analysis of mature ROs for rod photoreceptors (RHO), cone photoreceptors (PNA), photoreceptor connecting cilia (Rootletin), MG cells (SOX9) and MG projections (Vimentin) and F-actin. Scale bar: 10 µm. n = 3 ROs.

To validate proper retinal architecture, we conducted IF-analysis on w35 healthy-1 and USH1C-1 ROs. Both RO lines demonstrated the presence of rod (RHO) and cone (PNA) photoreceptor IS/OS, as well as the ciliary rootlet (Rootletin) (Fig. 1C). MG nuclei (SOX9) were observed in the ONL, with MG projections (Vimentin) spanning all retinal layers. Notably, examination of the OLM revealed that filamentous actin (F-actin) was less prominent in USH1C-1 ROs, which is consistent with the observed reduction in OLM thickness (Fig. 1B).

In summary, mature ROs demonstrated proper cellular organization after w35 of differentiation. However, USH1C-1 ROs displayed notable abnormalities. We observed OLM thinning from w30 and decreased amounts of F-actin in mature ROs, potentially indicating early OLM degeneration.

### Single-cell RNA sequencing showed changes in cellular composition and communication in USH1C-1 retinal organoids

To get an overview of the molecular pathology in USH1C, we compared the transcriptomes of ROs by performing 10X Genomics Chromium scRNA-seq. In total, we sequenced four healthy-1 and three USH1C-1 w35 ROs resulting in 20,020 single cell transcriptomes. After quality control, 19,173 cells remained, which were projected on a two-dimensional uniform manifold approximation and projection (UMAP) (Fig. S3A). Based on their marker gene expression (Table S4), 31 cell clusters were identified and assigned to a retinal cell type. We detected distinct types and maturation stages of RPCs, RPE cells, cone and rod photoreceptors, horizontal cells, bipolar cells, amacrine cells and MG cells, consistent with previous findings [25–27, 36, 61]. Interestingly, we also observed a small population of cells co-expressing MG and rod photoreceptor markers, which to our knowledge has not been reported in ROs before. We named this population Müller glia to rod cells (MGtR). From MGtR1 to MGtR3, cells showed a decrease in MG-specific gene expression, while the expression of rod photoreceptor-specific markers increased (Table S4).

To keep a better overview in subsequent analysis, we fused the 31 cell types based on similarities in their marker gene expression profile (Fig. S3B). In total, 18 different cell types remained (Fig. 2A) and their marker genes were rechecked (Fig. 2B; Table S5). We then performed pseudotime analysis using Monocle 3 to assess the temporal dynamics of gene expression during RO development. By setting early RPCs as a starting point, we observed a gradual transition in expression patterns from early RPCs to MG cells and to bipolar/photoreceptor cells (Fig. 2C). Mature rod photoreceptors marked the terminal and most differentiated state in pseudotime analysis.

**Fig. 2.**
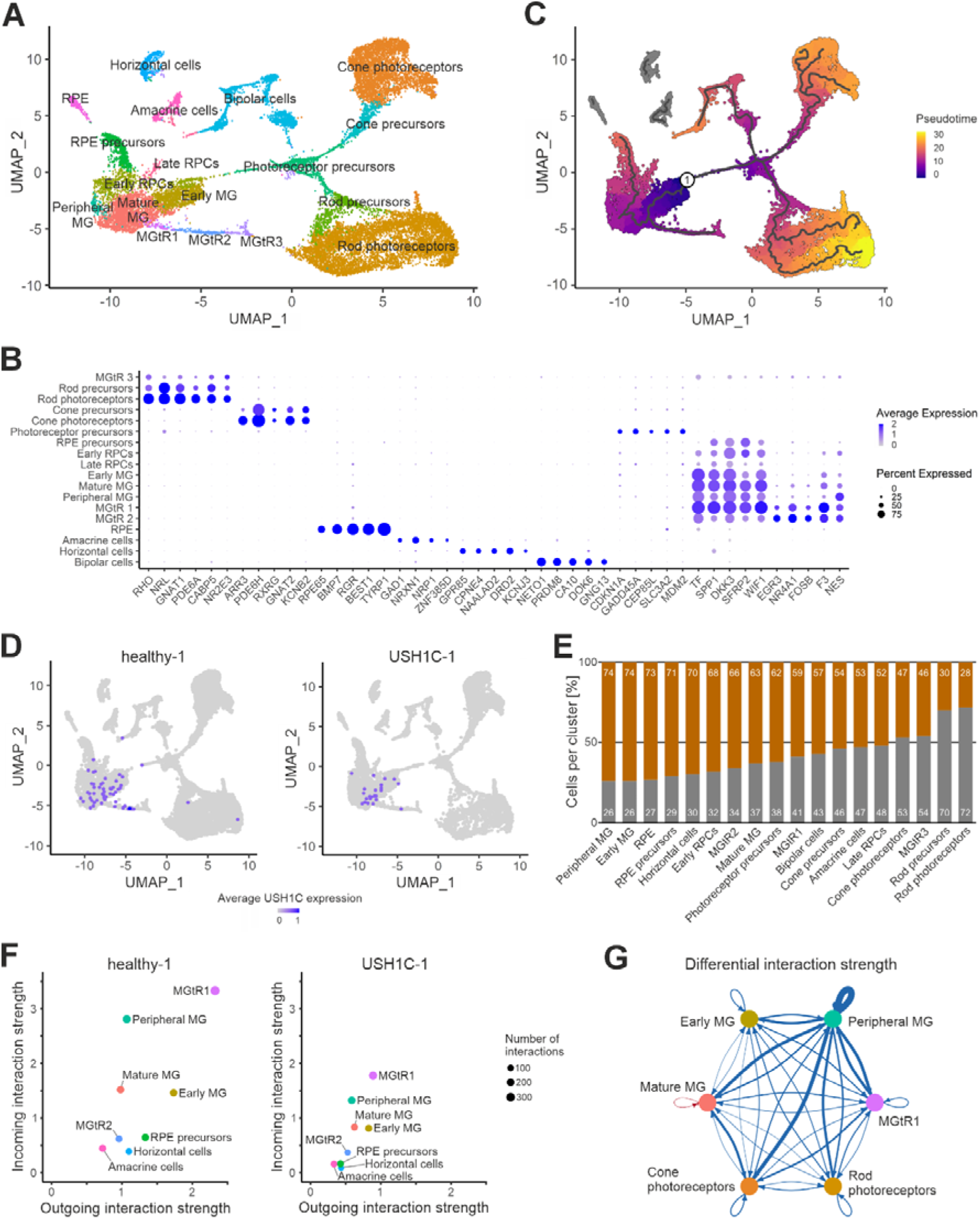
Single-cell RNA sequencing of mature healthy-1 and USH1C-1 retinal organoids. **A)** Two-dimensional Uniform Manifold Approximation and Projection (UMAP) of 18 different cell types based on the transcriptomes of 19,173 single cells. n = 4 healthy-1 and 3 USH1C-1 ROs. **B)** Expression of the top marker genes for each cell cluster. Blue color gradient indicates the average expression per cell cluster. Dot sizes indicate the portion of cells per cell cluster that express the corresponding marker. **C)** Pseudotime analysis displaying developmental trajectories of cells from an early undifferentiated state (blue) to a late differentiated state (yellow). The cluster of early RPCs (1) was manually set as the root node. **D)** UMAP of *USH1C* expression across the different cell types in healthy-1 and USH1C-1 ROs. Each blue dot corresponds to an *USH1C*-positive cell. Blue color gradient corresponds to the normalized *USH1C* expression per cell. **E)** Proportion of healthy-1 (dark grey) and USH1C-1 (dark orange) RO-derived cells for each cell type. **F)** Number of interactions and interaction strength for different cell types from healthy-1 and USH1C-1 ROs. Displayed are the eight most differential cell types of healthy-1 and USH1C-1 ROs. Outgoing interaction strength is shown on x-axis. Incoming interaction strength is shown on y-axis. The number of distinct interactions is indicated by dot size. **G)** Differential interaction strength between mature MG, early MG, peripheral MG, MGtR1, rod and cone photoreceptors. Blue arrows indicate a decreased interaction strength, red arrow indicates an increased interaction strength in USH1C-1 ROs compared to healthy-1 ROs. The thickness of the arrow correlates with the strength of the differential interaction.

To assess the expression pattern and levels of *USH1C* in ROs, we mapped *USH1C*-positive cells onto the UMAP. In healthy-1 ROs, only 0.5% of the total cell population expressed *USH1C* (Fig. 2D). USH1C-1 ROs revealed a minor difference, with 0.4% *USH1C*-positive cells. Both RO lines exhibited low *USH1C* expression levels in RPCs, early MG, mature MG and MGtR1/2, which is consistent with previous findings in mature ROs [25, 36]. Notably, we identified two additional *USH1C*-positive rod photoreceptor cells in healthy-1 ROs.

Despite the low expression of *USH1C* in both RO lines, we sought to investigate potential phenotypic alterations. Since Usher syndrome has been associated with degeneration and loss of photoreceptors [11, 62–64], we examined the number of healthy-1 and USH1C-1 cells across the different cell types to assess potential shifts in the cell population. Of 18 cell types, only five (cone precursors, amacrine cells, late RPCs, cone photoreceptors and MGtR3) showed relatively minor differences within 10% in the proportion of healthy-1 and USH1C-1 cells (Fig. 2E). Interestingly, USH1C-1 ROs exhibited a strong reduction in rod lineage cells, while cone photoreceptors remained unaffected. By contrast, various MG subtypes (peripheral MG, early MG, MGtR2, mature MG, MGtR1) were enriched in USH1C-1 ROs compared to healthy-1 ROs.

To determine potential alterations in cellular processes related to USH1C, we used CellChat for inferring intercellular communication from our scRNA-seq data [65]. For healthy-1 and USH1C-1 ROs, CellChat quantified the diversity and expression levels of ligand-receptor pairs and thus assessed the number and strength of intercellular interactions (Fig. S3C). Comparison of both RO lines revealed that the total number of interactions decreased only slightly in USH1C-1 ROs (Fig. S3D). However, the overall interaction strength was reduced by almost half, suggesting that communication efficiency, rather than communication diversity, is compromised in USH1C-1 ROs (Fig. S3E). We decided to examine the differences in interaction strength and assessed the most communicative cell types. In both RO lines, peripheral MG, MGtR1, mature MG and early MG cells had the highest incoming and one of the highest outgoing interaction strengths, indicating their key role in intercellular communication (Fig. 2F, Fig. S3F). Interestingly, comparative analysis revealed that signaling of these four MG cell types was most reduced in USH1C-1 compared to healthy-1 ROs (Fig. 2F; Table S6, S7). When specifically evaluating the interaction strength between MG and photoreceptor cells, we observed a strong downregulation of communication efficiency in USH1C ROs, indicating reduced information exchange between these cell types (Fig. 2G).

Overall, *USH1C* expression in mature ROs was found to be low and predominantly restricted to MG cells. We observed a reduction in the number of rod photoreceptors accompanied by an increase in MG cells. Moreover, intercellular communication efficiency was decreased in USH1C-1 ROs, with MG subtypes being particularly affected.

### Single-cell RNA sequencing revealed Müller glia and photoreceptor pathology in USH1C-1 retinal organoids

To elucidate the signaling pathways targeted by intercellular communication in healthy-1 and USH1C-1 ROs, we employed CellChat to examine the communication patterns for all cell types. A comparison of outgoing and incoming signaling pathways across all communication patterns revealed no substantial differences, suggesting that cells of both RO lines sent and receive similar signals (Fig. 3A; Fig. S4A). Moreover, the diversity of signaling pathways remained largely similar between healthy-1 and USH1C-1 for both outgoing (Fig. 3A) and incoming (Fig. S4A) patterns, which is consistent with previous analysis (Fig. S3D). Notably, we observed that MG subtypes (peripheral MG, MGtR1, mature MG and early MG cells) were predominantly grouped within pattern 1 in healthy and USH1C-1 ROs, indicating similar signaling profiles in these cells. Since MG cells are responsible for most of the communication, we investigated their outgoing and incoming signaling pathways in more detail. As expected, we found consistent signaling across the different MG subtypes, which was mainly related to retinal adhesion (Fig. 3B-E; Fig. S4B-E). Specifically, we observed signaling pathways associated with extracellular matrix (ECM)-receptor interactions (collagen, FN1, laminin, RELN, SPP1), focal adhesions (RELN, SPP1), cell adhesions (CADM, CDH, CD99, CNTN, FLRT, NCAM, NECTIN, NRXN, PCDH, PTRP, SLITRK) and adherens junctions (CDH, NECTIN, PTRP) (Table S8). Strikingly, the majority of these signaling pathways was significantly downregulated in USH1C-1 ROs.

**Fig. 3.**
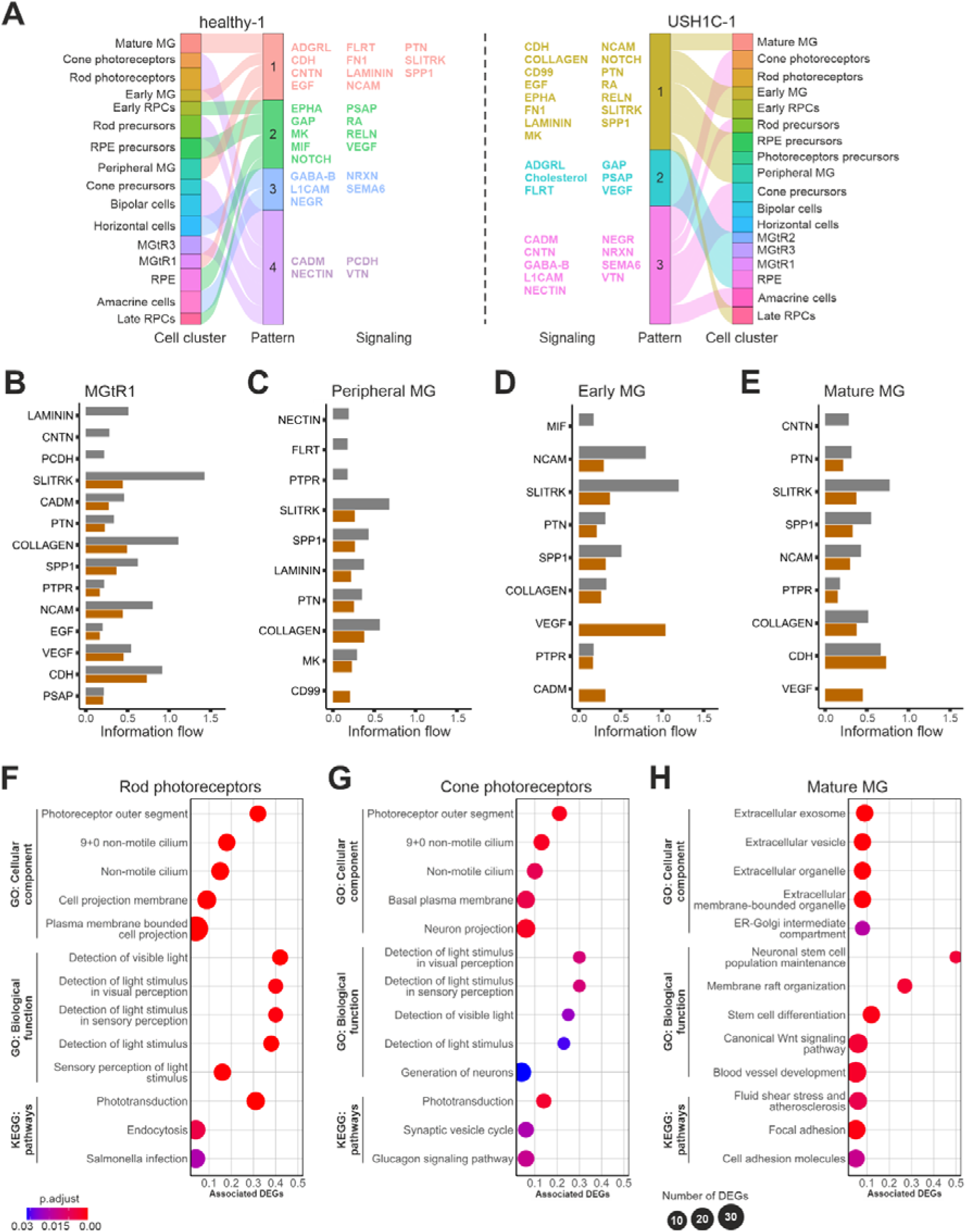
Outgoing intercellular signaling and differential expressed gene analysis in mature healthy-1 and USH1C-1 retinal organoids. **A)** River plot grouping cell types of healthy-1 and USH1C-1 ROs according to similarities in their outgoing communication. Each signaling pattern is defined by different signaling pathways. **B-E)** Comparison of significant differential outgoing signaling pathways from MGtR1, peripheral MG, early MG and mature MG cells of healthy-1 (dark grey) and USH1C-1 (dark orange) ROs. Statistical analysis was done using Wilcoxon test. **F-H)** GO-term and KEGG analysis of differential expressed genes (DEGs) in rod photoreceptors, cone photoreceptors and mature MG cells. Displayed are the top five GO-terms for Cellular component and Biological function and top three KEGG pathways terms (according to p.adjust). p.adjust is indicated by color code. Number of identified DEGs related to a term is indicated by dot size. X-axis defines the proportion of associated DEGs, i.e. the number of DEGs normalized to the total number of genes assigned to a specific GO-/KEGG-term.

To further elucidate the underlying mechanisms and downstream effects of altered communication in USH1C, we performed DEG analysis between healthy-1 and USH1C-1 ROs across all cell types (Table S9). Rod photoreceptors (253 DEGs), cone photoreceptors (233 DEGs) and mature MG cells (192 DEGs) were identified as the most affected cell types in USH1C-1 ROs (Table S10). We decided to conduct GO and KEGG analysis on the cells (Table S11) and summarized the top five GO- and top three KEGG-terms. In rod photoreceptors, GO analysis of DEGs revealed enrichment for Cellular Component terms related to the photoreceptor outer segment and non-motile cilium in USH1C-1 ROs (Fig. 3F). Moreover, Biological Process terms associated with light detection were enriched. KEGG pathway analysis further demonstrated a DEG enrichment in phototransduction. Notably, we observed nearly identical GO- and KEGG-terms when analyzing cone photoreceptors (Fig. 3G). Manual literature research for both rod and cone photoreceptors additionally revealed that even more genes related to phototransduction were dysregulated in USH1C-1 than those detected by automated KEGG analysis (Table S12) [66]. When looking at mature MG, GO- and KEGG-terms appeared to be more diverse (Fig. 3H). GO analysis of DEGs showed enrichment for Cellular Component terms associated with the extracellular exosome in USH1C-1 ROs. Since extracellular vesicles serve as important mediators of intercellular signaling by transporting DNA, RNA and proteins between cells [67], these results support our CellChat analysis of impaired communication efficiency in USH1C. Additionally, Biological Process terms revealed a DEG enrichment in canonical Wnt signaling, which we recently showed to be dysregulated in USH1C [23]. KEGG pathway analysis lastly indicated DEGs to be enriched in focal adhesion and cell adhesion molecules being in line with the downregulation of these signaling pathways in our CellChat analysis.

In summary, our findings in ROs highlighted MG cells as major drivers of intercellular communication, primarily exchanging signals related to retinal adhesion. While this signaling was prominent in both RO lines, its efficiency was downregulated in USH1C-1. Coinciding with that, we observed DEGs associated with IS/OS development and function in rod and cone photoreceptors, marking a potential consequence of altered intercellular communication.

### Phototransduction is partially impaired in USH1C retinal organoids

To enhance the robustness of analyses, we decided to expand our cellular model. We used USH1C fibroblasts from a sibling of the USH1C-1 donor carrying identical pathogenic variants and reprogrammed them into hiPSCs (USH1C-2) (Fig. S1A). Characterization of USH1C-2 hiPSCs and an additional control line (healthy-2) verified the presence and absence of the pathogenic variant, respectively, confirmed genetic stability, and demonstrated typical stem cell features (Fig. S5A-E). We differentiated the two lines into ROs and repeated our morphological characterization. Notably, USH1C-2 ROs showed a significant reduction in photoreceptor IS/OS length from w30 to w35 compared to all other RO lines (Fig. S6A). Consistent with our observations in USH1C-1 ROs, the OLM thickness in USH1C-2 ROs decreased from w30 to w35. Additionally, we observed less rod (RHO) and cone (PNA) photoreceptors, as well as mislocalized MG cells (SOX9) and no OLM-associated F-actin (Phalloidin) in USH1C-2 ROs, indicating a more severe pathology than in USH1C-1 ROs (Fig. S6B).

To further explore the transcriptional and morphological impairments associated with photoreceptors in USH1C-1 and USH1C-2 ROs, we conducted IF-analysis of different phototransduction proteins. We processed the obtained images in Fiji by applying a default threshold and quantified the protein-positive area apical to the ONL for rod (RHO, GNAT1, GUCA1A, SAG) and cone photoreceptors (GUCA1A, ARR3) (Fig. 4A). Interestingly, we observed no difference in the size of phototransduction protein-positive IS/OS regions among healthy-1, healthy-2, and USH1C-1 ROs. In contrast, USH1C-2 ROs exhibited a significant reduction in phototransduction protein-positive IS/OS regions, consistent with the observed IS/OS reduction in our morphological analysis (Fig. S6A, S6B).

**Fig. 4.**
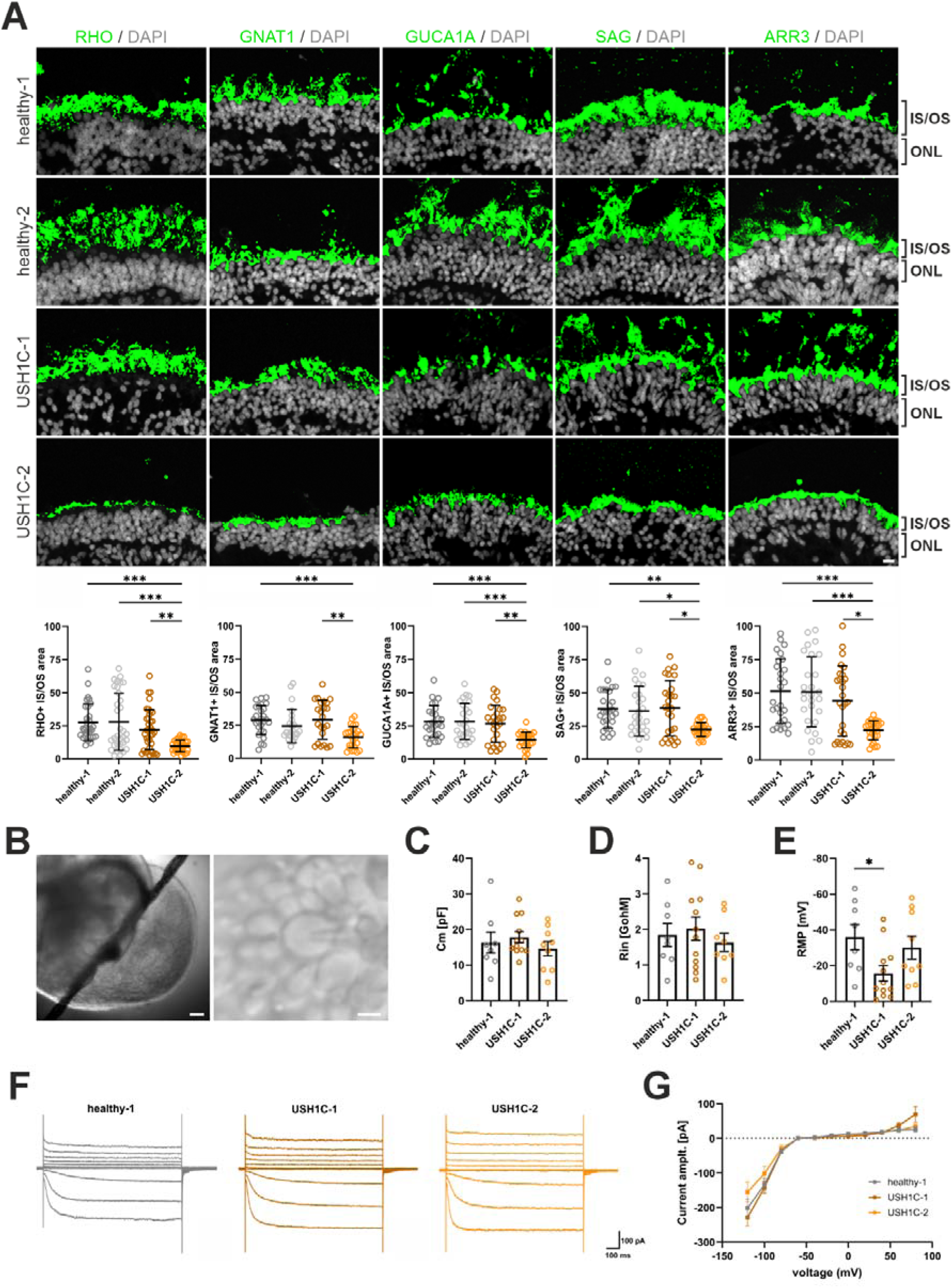
Morphological and electrophysiological characterization of mature photoreceptor inner and outer segments. **A)** Representative images processed for IF-quantification of phototransduction proteins in mature healthy-1, healthy-2, USH1C-1 and USH1C-2 ROs. Scale bar: 10 µm. Phototransduction proteins Rhodopsin (RHO; rods), Opsin (cones), G protein subunit alpha transducin 1 (GNAT1: rods), Guanylate cyclase activator 1A (GUCA1A: rods and cones), S-antigen visual arrestin (SAG: rods) and Arrestin-3 (ARR3: cones) are marked. Kruskal-Wallis test for n = 3 ROs per cell line, 2-4 sections per organoid, 2-4 random images per section. Each data point corresponds to one image. **B)** Representative RO located in the recording chamber for electrophysiological recordings (left picture). A smooth, round cone photoreceptor OS with the patch-clamp electrode attached (right picture). Scale bar: 100 µm and 10 µm. Diagrams showing **C)** the mean membrane capacitance (Cm), **D)** the mean input resistance (Rin) and **E)** the mean resting membrane potential (RMP). One-way ANOVA on n ≥ 3 ROs per cell line, ≥ 2 cells per RO. Each data point corresponds to one cell. **F)** Voltage-clamp recordings showing the membrane current of representative ROs at different holding potentials from -120 mV to +80 mV in 20 mV increments for healthy-1, USH1C-1 and USH1C-2 ROs. Scale bar: 100 ms and 50 pA. **G)** Mean current-voltage-relationships (I-V curves) from healthy-1, USH1C-1 and USH1C-2 ROs. Two-way ANOVA on n ≥ 3 ROs per cell line, ≥ 2 cells per RO. Each data point corresponds to one cell.

Next, we investigated potential changes in the electrical membrane properties of photoreceptors in USH1C by performing whole-cell patch-clamp recordings in cone photoreceptors. For recordings, only smooth and bright cone OS without signs of degeneration were selected from healthy-1, USH1C-1 and USH1C-2 ROs (Fig. 4B). Measurements of the membrane capacitance (Cm) and input resistance (Rin) revealed no differences among the RO lines, indicating comparable cell membrane properties between photoreceptors (Fig. 4C, 4D). We then assessed the resting membrane potential (RMP), which ranged from -2 to -65 mV (Fig. 4E). This was consistent with previous results using the same protocol [31]. Interestingly, USH1C-1 cones were significantly more depolarized than healthy-1 cones (-16 mV vs. -36 mV). USH1C-2 cones showed a tendency towards a more depolarized state (-30 mV), although this difference was not significant. Lastly, we investigated the current-voltage (I-V) relationship in our patch-clamp recorded ROs. Here, we applied gradually depolarizing voltage steps (+20 mV) from -120 mV to +80 mV to calculate the corresponding I-V curve for each cell (Fig. 4F, 4G). We did not observe any changes in these I-V curves between ROs. Since hyperpolarization-activated cyclic nucleotide-gated (HCN) channels and cyclic nucleotide-gated (CNG) channels are known to mediate the I-V curve in ROs [31] we conclude that the functional properties of HCN-channels as well as of the CNG-channels are likely not altered between healthy-1 and USH1C cone photoreceptors. These results align with our scRNA-seq analysis, which revealed no transcriptional differences for HCN and CNG channels in USH1C-1 cone photoreceptors (Table S2).

In conclusion, these data confirm a photoreceptor pathology in USH1C-1 and USH1C-2 ROs, albeit to different degrees. While photoreceptor IS/OS were diminished in USH1C-2 ROs, USH1C-1 ROs showed no signs of IS/OS reduction, but functional impairments in cone photoreceptors.

### USH1C retinal organoids show an impaired outer limiting membrane architecture

Previous studies demonstrated that MG cells can undergo activation and proliferation upon retinal degeneration [68, 69]. To see whether alterations in photoreceptor morphology were accompanied by changes in MG cell behaviour, we conducted IF-analysis to quantify the number of SOX9-positive MG cells (Fig. 5A). USH1C-1 and USH1C-2 ROs exhibited an increase in the MG cell population compared to healthy-1 ROs (Fig. 5B), consistent with our scRNA-seq results (Fig. 2E). However, MG cell numbers were also elevated in healthy-2 ROs, indicating that an increase in MG cells might not be exclusively linked to pathological processes. As migration of MG cells has been found as an additional marker of retinal disease [70], we analyzed the amount of mislocalized SOX9-positive MG cells to the NRL-positive ONL. Quantification revealed an increased migration of SOX9-positive cells in USH1C-2 ROs, implying a compensatory response (Fig. 5C) [71].

**Fig. 5.**
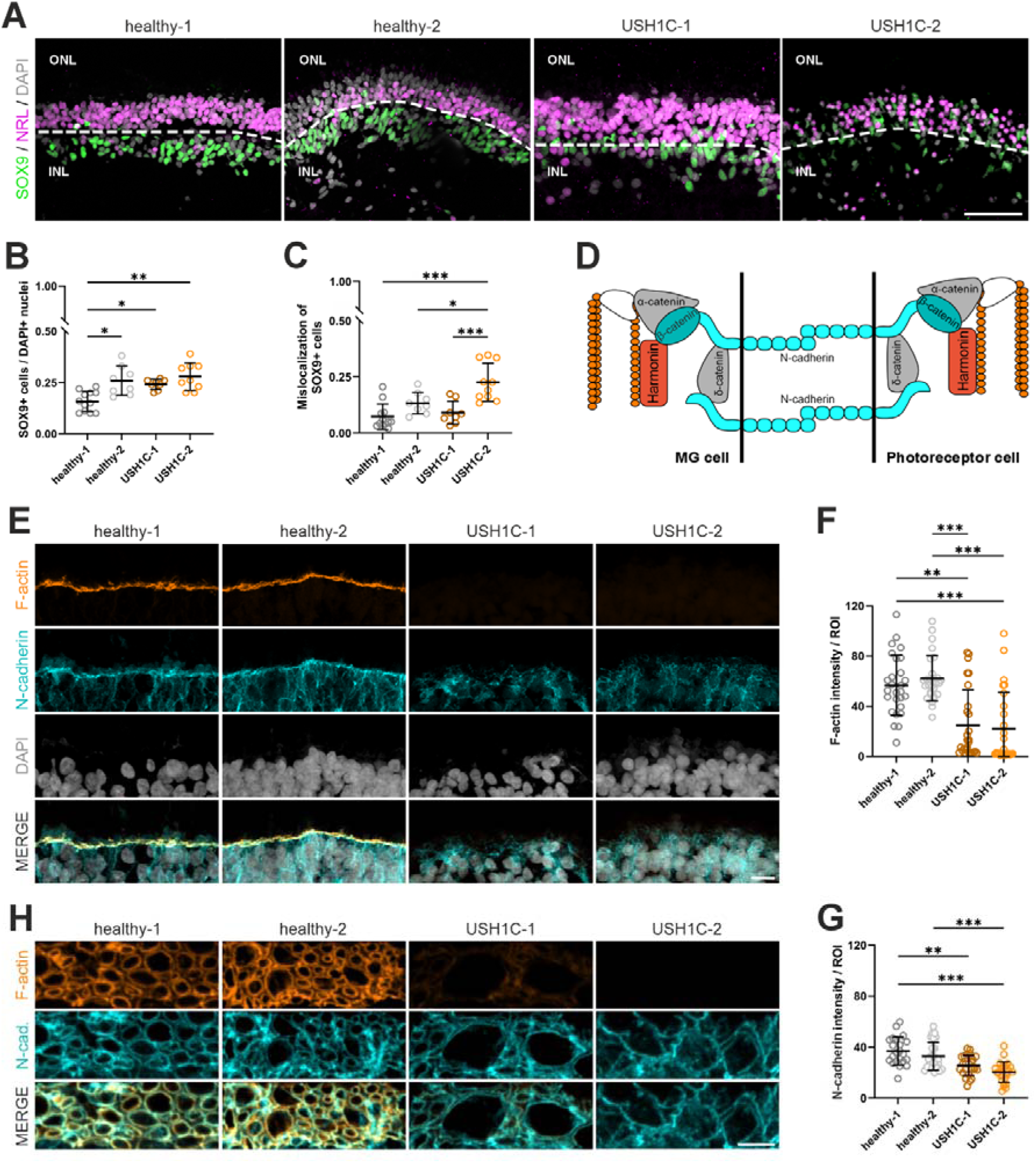
Morphological analysis of the outer limiting membrane and Müller glia cells in mature retinal organoids. **A)** Representative images showing SOX9-positive MG nuclei. The outer nuclear layer (ONL) was assessed via NRL-positive nuclei. The inner nuclear layer (INL) was determined via SOX9-positive cells. Scale bar: 50 µm. **B-C)** Total number of MG cells and number of mislocalized MG cells to the ONL. Statistical analysis was done using Kruskal-Wallis test for **B)** and One-way ANOVA for **C)** on n = 3 ROs per cell line, 3 entire sections per RO. Each data point corresponds to one entire section. **D)** Schematic of the OLM-related adherens junction between MG and photoreceptor cells. **E)** IF-analysis of OLM-related F-actin and N-cadherin. Scale bar: 10 µm. **F-G)** Quantification of OLM-related F-actin and N-cadherin intensities. Kruskal-Wallis test for n = 3 ROs per cell line, 3 sections per organoid, 3 random images per section, 10 random regions of interest (ROIs) per image. Each data point corresponds to one image. **H)** IF-analysis of OLM-related F-actin and β-catenin. Scale bar: 5 µm.

Our transcriptomic analyses revealed dysregulation related to retinal adhesion in MG cells of USH1C-1 ROs (Fig. 3B-E, 3H; Fig. S4B-E). Moreover, our initial morphological characterization provided evidence of impaired OLM development and morphology in both USH1C-1 and USH1C-2 ROs (Fig. 1B, 1C; Fig. S6A, S6B). We therefore quantified the OLM by analyzing the fluorescence intensity of F-actin and N-cadherin apical to the ONL. Together with catenins and harmonin [4, 72], both components are crucial mediators of the OLM-related adherens junction between MG and photoreceptor cells (Fig. 5D). Strikingly, USH1C ROs showed a strong decrease and in some cases, a complete absence of F-actin (Fig. 5E, 5F). Additionally, N-cadherin levels were reduced in USH1C ROs, resulting in a less prominent OLM structure (Fig. 5E, 5G). Super-resolution microscopy of F-actin and β-catenin further revealed a less organized OLM architecture leading to larger pores in both USH1C RO lines (Fig. 5H).

Overall, our findings demonstrate that USH1C ROs exhibit a disrupted OLM architecture accompanied by partially altered MG cell behavior. Together with the altered communication observed in our scRNA-seq, these results indicate that impaired adhesion mechanisms contribute to compromised retinal integrity in USH1C.

## Discussion

Rodent models are widely used to advance our understanding of neurodegenerative diseases [73]. However, in the case of USH1C, they fail to recapitulate retinal pathology observed in patients [1, 9, 10]. Recent studies have demonstrated the utility of patient-derived cellular models for unraveling retinal disease mechanisms, offering a promising alternative to animal studies [33–35]. Here, we established the first USH1C RO model, generated from hiPSCs of two siblings, allowing systematic exploration of the molecular processes underlying retinal degeneration. Our ROs recapitulated various retinal cell types, including photoreceptor and MG cells, in which harmonin is predominantly expressed [4, 11, 12]. Our findings revealed a complex pathology affecting both cell populations, manifesting in a disrupted OLM architecture and dysregulated intercellular communication.

### ROs recapitulate patient-specific variations in photoreceptor pathology

Retinal pathology in Usher syndrome is characterized by an impaired architecture and progressive degeneration of photoreceptors [11, 74–76]. Recent studies have highlighted harmonin’s role in maintaining photoreceptor structure and function. Specifically, harmonin localizes within rod OS at the disc-forming base and at the cytoplasmic side of discs membranes, suggesting two potential functions [4]. First, harmonin appears to be involved in disc stacking as pathogenic variants were associated with disrupted disc architecture [11]. And second, harmonin may indirectly influence phototransduction by modulating RHO organization or function through interaction with it [4]. Although we observed nearly no *USH1C* expression in RO photoreceptors, pathogenic variants led to altered IS/OS development and transcriptional, morphological and functional impacts on phototransduction, underscoring harmonin’s importance for photoreceptor viability. Notably, we observed substantial differences in photoreceptor pathology between USH1C ROs, with USH1C-2 ROs showing a more severe phenotype compared to USH1C-1 ROs. These results align with optic coherence tomography (OCT) data from the patient’s eyes, mirroring a stronger disease progression for the USH1C-2 donor [4]. We assume that donor-specific genetic background and epigenetic regulation contribute to these variations in the spatiotemporal dynamics of photoreceptor pathology [77, 78]. Moreover, our study highlights the capability of ROs to model inter-patient variability.

### Loss of OLM architecture as a common disease marker

The OLM is a network of heterotypic adherens junctions between the apical processes of MG and the IS of photoreceptor cells [79]. Within these junctions, harmonin functions as a molecular linker, connecting cadherin-catenin complexes to the intercellular actin cytoskeleton through its interactions with β-catenin and the actin-binding protein filamin A [4, 23]. Additionally, the harmonin b isoform directly binds F-actin via its PST domain [80]. Our morphological analyses revealed OLM thinning during RO development, which coincided with reduced F-actin and N-cadherin levels at the OLM in both mature USH lines. Recent *in vitro* studies have demonstrated bi-directional coupling of cadherin and F-actin dynamics at adherens junctions, showing that cadherin pathogenic variants disrupt these dynamics [81]. Given harmonin’s role at the interface between N-cadherin and F-actin, we suggest that pathogenic variants of harmonin similarly destabilize adherens junction complexes. Supporting this hypothesis, we showed that USH1C ROs exhibited an altered OLM architecture characterized by larger pores. Dysregulation of the OLM architecture has been previously linked to multiple inherited retinal diseases [35, 85, 88], underscoring its importance for retinal homeostasis. Remarkably, pathogenic variants of clarin-1, an interaction partner of harmonin causing USH3A, lead to comparable OLM degeneration and reduced N-cadherin levels [83]. Together, these findings emphasize the essential role of harmonin and other USH proteins in maintaining OLM integrity, and highlight the OLM as a sensitive and promising marker for evaluating preclinical therapeutic strategies.

### MG-mediated intercellular communication is essential for maintaining retinal integrity

Intercellular communication plays a fundamental role in coordinating cellular development, maintenance, and function through the exchange of signals. In complex tissues like the retina, elucidating these cellular interactions is critical for understanding the mechanisms that preserve tissue integrity and homeostasis [84]. Dysregulation of intercellular communication has been implicated in the pathogenesis of various retinal diseases [85–87]. Our CellChat analysis identified MG cells as the most active signaling population in ROs, highlighting their role as key drivers of retinal communication [88, 89]. Signaling activity of MG cells was predominantly mediated by physical, contact-dependent communication, involving cell-cell adhesions (adherens junctions, cell-cell adhesion molecules) and cell-ECM adhesions (ECM-receptor interaction, focal adhesions). These findings are consistent with the established role of MG cells in providing mechanical stability and structural support in the retina, as they span all retinal layers and maintain contact with all retinal neuron [90, 91]. Importantly, USH1C-1 ROs revealed that MG-mediated communication through these adhesion pathways is severely impaired in both directions between MG and retinal neurons. To date, harmonin has been shown to primarily participate in adherens and tight junctions at the OLM [4], and our data provide evidence that harmonin dysfunction leads to OLM degeneration. We thus propose that the observed defects in other contact sites arise as secondary consequences of the primary OLM disruption. Although such effects have not been studied in the retina, analysis of other tissues demonstrates a dependence of cell-cell and cell-ECM adhesions, where disruption of one adhesion system can perturb the other, potentially via alterations in the actin cytoskeleton [92, 93]. Consistent with this, we observed aberrant F-actin organization at the OLM and transcriptional dysregulation of ECM components [90], such as collagen, fibronectin and laminin in CellChat and DEG analysis. Given that MG cells are the principal producers of ECM components in the retina, their imbalance in the expression of these components likely contributes to an overall impairment of retinal integrity. Beyond MG cells, other retinal cell types such as bipolar cells, cone photoreceptors, early RPCs and RPE precursors exhibited DEGs related to physical cell contacts, suggesting a more widespread failure in maintaining retinal structure.

### The dual role of harmonin and **β**-catenin in retinal homeostasis

In the retina, β-catenin serves a dual function in tissue morphogenesis and intracellular signaling. During morphogenesis, β-catenin is essential for the formation and maintenance of adherens junction complexes, thereby ensuring proper tissue organization. As a signaling molecule, β-catenin functions as key transcription factor in the canonical Wnt signaling pathway, regulating the expression of genes involved in tissue development and differentiation [94]. We have previously shown that harmonin directly interacts with β-catenin to modulate canonical Wnt signaling, and that both proteins co-localize at the OLM [4, 23]. Our current findings in ROs further highlight harmonin’s regulatory effect on β-catenin in both structural and signaling context. Notably, we observed prominent β-catenin staining at the OLM, which coincided with a disrupted OLM architecture in USH1C ROs. Additionally, transcriptomic analyses revealed an enrichment of various DEGs associated with canonical Wnt signaling in early RPCs and mature MG. We hypothesize that the disruption of the harmonin/β-catenin-dependent adherens junctions not only perturbs the structural integrity of the OLM and other contact sites but also affects intracellular canonical Wnt signaling, thereby triggering MG cell-specific responses to retinal injury [95–97]. Consistent with previous studies describing MG proliferation and migration as injury responses [98–100], our transcriptomic and morphological analyses revealed increased numbers of MG cells and their migration into the ONL, suggesting the activation of compensatory mechanisms. Although our RO model did not exhibit GFAP expression, as confirmed by scRNA-seq and IF-analysis (data not shown), we observed elevated *VIM* expression in MG cells. This upregulation may reflect an initial reactive gliosis-like response, potentially serving to stabilize the retinal architecture in the face of OLM disruption, compensating for the loss of structural integrity [99].

### Human MG cells have a regenerative potential

Currently, no treatment exists for USH1C-related retinal degeneration. While the regenerative potential of MG cells is promising for visual restoration, their capacity varies significantly between species [101]. In non-mammalian species, MG cells are able to acquire retinal stem cells characteristics, generating new neurons post-injury [102–106]. In mammals, however, MG cell regeneration is limited and requires molecular intervention to enhance efficacy [107]. Using scRNA-seq, we identified three MGtR cell types exhibiting shifts from MG-specific to rod photoreceptor-specific gene expression. While these cell populations have not yet been identified in ROs, a study on RP mice reported MG subpopulations with elevated photoreceptor maintenance gene expression, suggesting a compensatory response [108]. Importantly, our data indicate that MGtR emergence is not exclusive to USH1C condition, implying a conserved re-differentiation mechanism rather than a disease-specific response. Nevertheless, MGtR cells of USH1C-1 ROs exhibited altered expression of genes involved in retinal adhesion and phototransduction. These findings suggest that *USH1C* pathogenic variants may impair the efficiency of MG re-differentiation, potentially limiting their regenerative capacity.

## Conclusion

In conclusion, our study provided novel insights into the molecular and cellular mechanisms underlying USH1C-related retinal degeneration using the first patient-derived USH1C RO model. We revealed that harmonin dysfunction compromises photoreceptor and MG cells, leading to a loss of OLM integrity and impaired intercellular communication. These defects propagate through the retinal tissue, compromising structural stability and signaling pathways essential for retinal homeostasis. By elucidating these aspects of disease progression, our work advances the understanding of USH1C pathology and provides valuable molecular and cellular readouts for future therapeutic development.

## Acknowledgements

First, we would like to thank the patients whose cell donations made this study possible. We appreciate Vasiliki Kalatzis, Carla Sanjurjo-Soriano and Carla Jimenez (University of Montpellier) for their hands-on training in RO generation. We also thank Pablo Llavona Juez (IMB Genomics Core Facility) for his assistance with scRNA-seq library preparation and sequencing. We acknowledge the Light Microscopy Core Facility (JGU Mainz) for access to the ZEISS LSM 900, funded by the Ministry of Science and Health of Rhineland-Palatinate and the European Regional Development Fund (ERDF/REACT-EU, Grant No. 84012490). Finally, we thank Joshua Linnert, Ulrike Maas and Yvonne Kerner (JGU Mainz) for their technical expertise, and Uwe Wolfrum (JGU Mainz) for scientific discussions.

## Statements and Declarations

### Author contributions

This study was designed, coordinated and supervised by NW and KNW. Human dermal fibroblasts were obtained by PDA, KS and SK. hiPSCs were obtained by DA, KZ, MEC and NW. hiPSCs were characterized by NW and KH. ROs were generated and characterized by NW. scRNA-seq was carried out by NW and MML. Quality control of scRNA-seq raw data and analysis consulting were performed by SiK. Analysis of scRNA-seq data was performed by MZ and NW. SOX9-positive cells were quantified by SeK, CR and NW. Photoreceptor IS/OS measurements were assessed by NW. Patch clamp experiments were performed by QW and supervised by TM. OLM analyses were done by NW. The manuscript was written by NW. All authors reviewed and approved the final version of the manuscript.

### Data availability

scRNA-seq data generated during the current study are available at NCBI Gene Expression Omnibus database as (publicly available after manuscript acceptance). All original code for analysis of scRNA-seq data, cell counting and intensity measurements has been deposited at://github.com/LabWolfrum/USH1C_retinal_organoids_Wenck/tree/main. Any additional information required to reanalyze the data reported in this paper is available from the lead contact upon request.

### Ethics approval

This study was performed in line with the principles of the Declaration of Helsinki. Procedures involving human material were ethically approved by the State Medical Association Rhineland-Palatinate (2021-16167).

### Consent to publish

The authors affirm that human research participants provided informed consent for publication.

