## Supplemental Information for "Lost in communication: How Müller glia cells fail to maintain retinal integrity in USH1C retinal organoids"

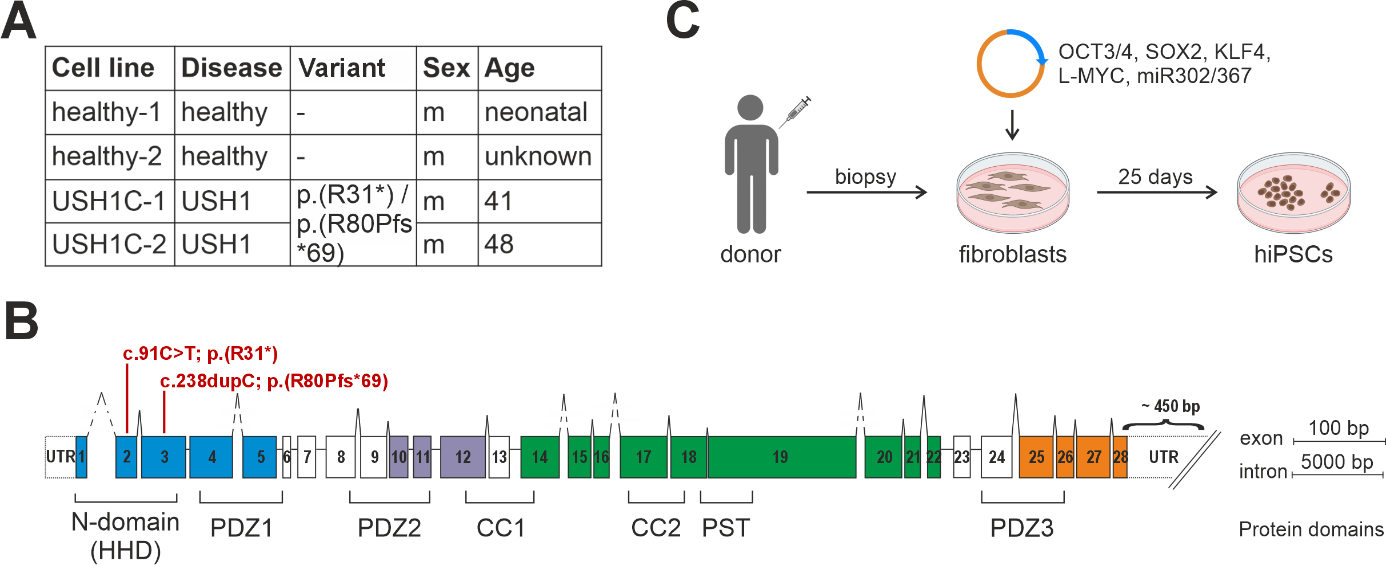

**Fig. S1 Characteristics of human dermal fibroblasts. A)** Fibroblast lines with corresponding disease state of the donor, pathogenic variants, sex and age at biopsy. **B)** Schematic of the *USH1C* gene. The *USH1C* gene consists of 28 exons that can form up to seven different protein domains. Patient-specific pathogenic variants c.91C>T; p.(R31*) and c.238dupC; p.(R80Pfs*69) are indicated in exon 2 and 3. HHD (harmonin homology domain), PDZ (PSD-95, DLG, ZO-1) domain, PST (proline–serine–threonine rich) domain, CC (coiled-coil) domain. **C)** hiPSCs were generated from donor fibroblasts by episomal reprogramming using the Yamanaka factors *OCT3/4*, *SOX2*, *KLF4*, *L-MYC* and miR302/367.

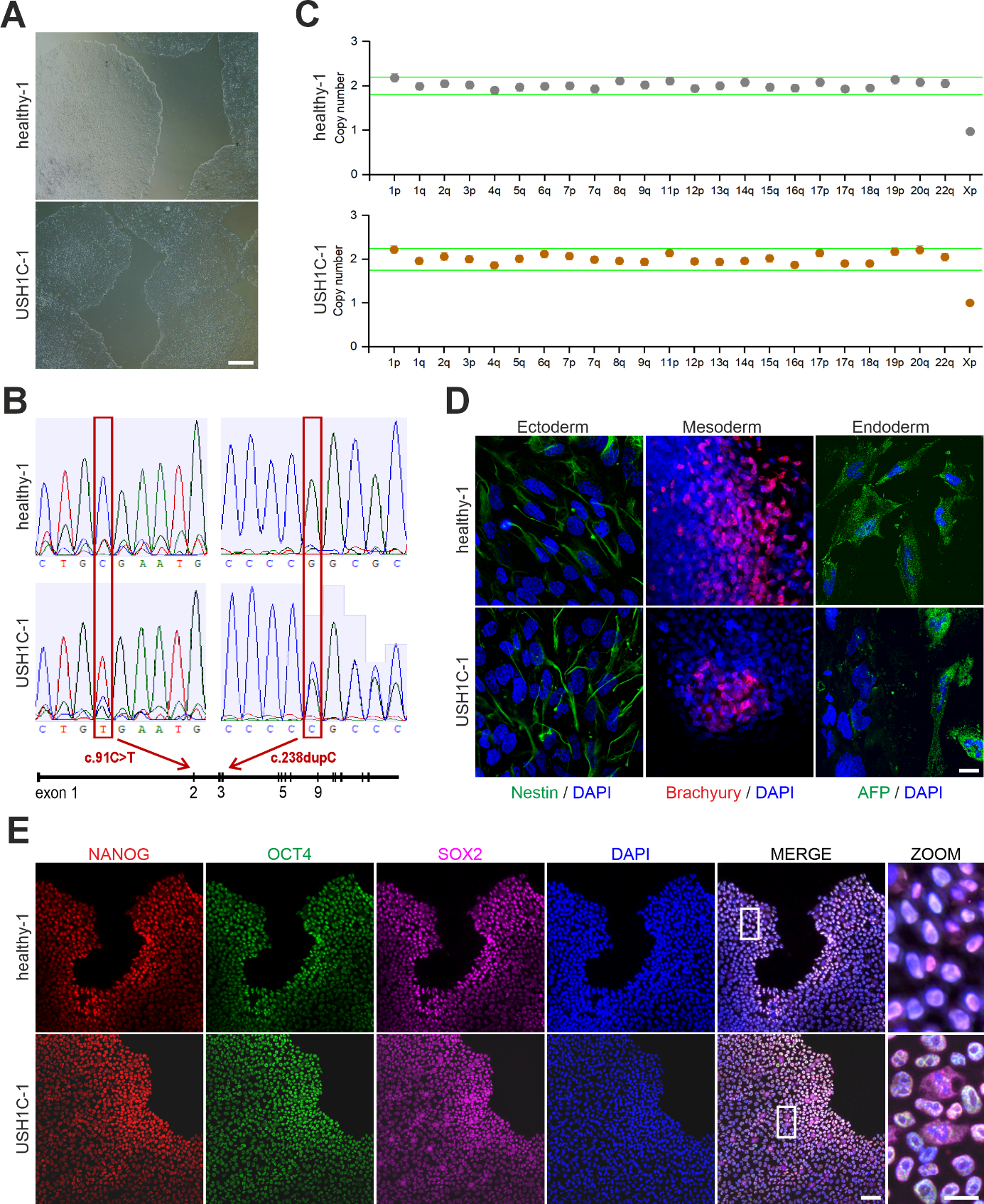

**Fig. S2 Characterization of healthy-1 and USH1C-1 human induced pluripotent stem cells. A)** Morphology of healthy-1 and USH1C-1 human induced pluripotent stem cell (hiPSC) colonies. Scale bar: 500 µm. **B)** Sanger sequencing determined the absence or presence of the c.91C>T; p.(R31*) and the c.238dupC; p.(R80Pfs*69) *USH1C* pathogenic variants in exon 2 and 3, respectively. **C)** Genomic stability of healthy-1 and USH1C-1 hiPSCs was assessed by testing for copy number variations (CNVs). **D)** Differentiation ability of healthy-1 and USH1C-1 hiPSCs into the three germ layers was assessed by an embryoid body (EB)-assay. Nestin was used as an ectodermal, Brachyury as a mesodermal and AFP as an endodermal marker. Scale bar: 25 µm. **E)** Expression of transcription factors NANOG, OCT4 and SOX2 confirmed the undifferentiated state of healthy-1 and USH1C-1 hiPSCs. Scale bars: 75 µm and 25 µm.

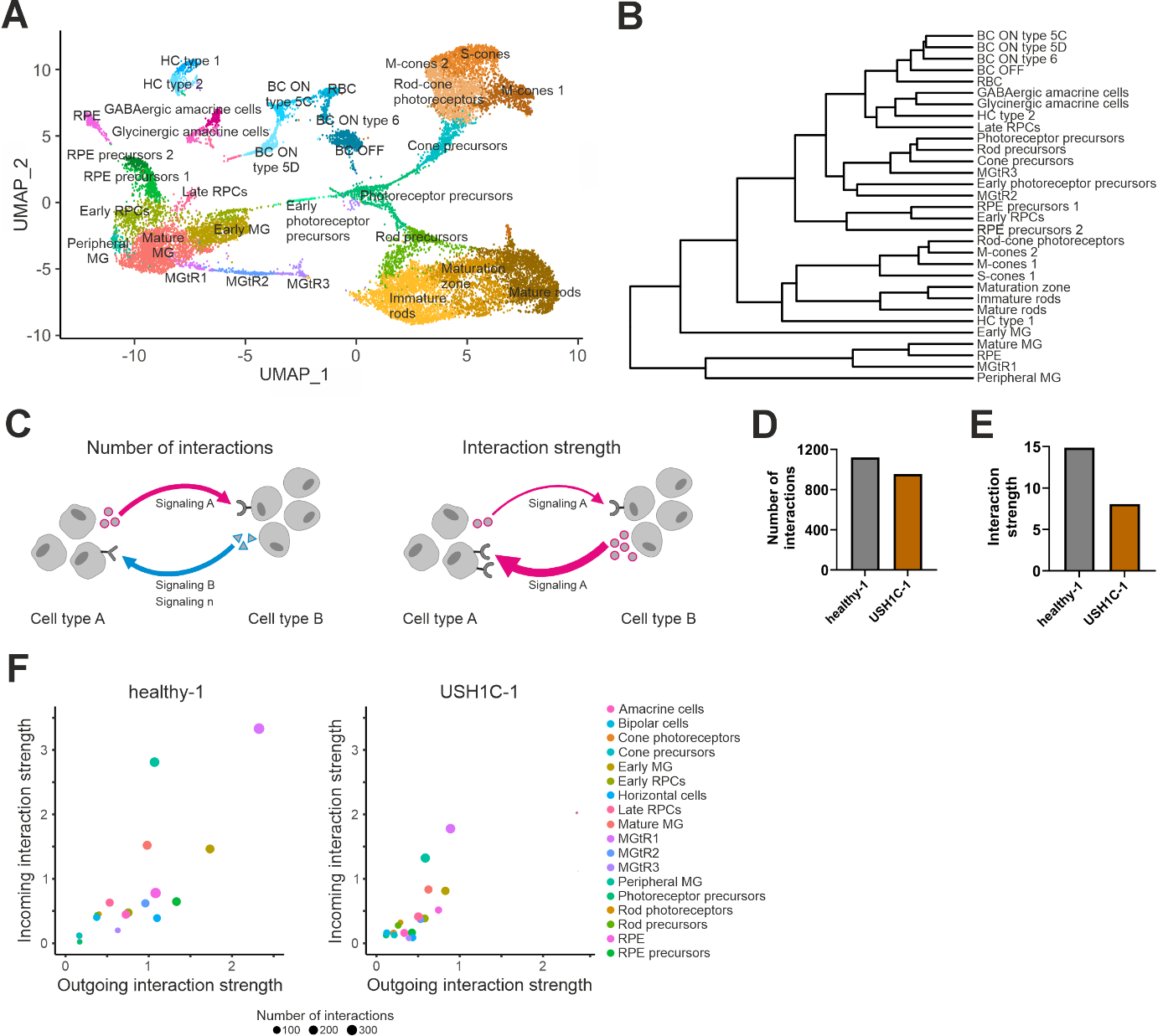

**Fig. S3 Single-cell RNA sequencing of mature healthy-1 and USH1C-1 retinal organoids. A)** Two-dimensional UMAP representation of the 31 different cell types identified based on the transcriptomes of 19,173 single cells. n = 4 healthy-1 and n = 3 USH1C-1 ROs. **B)** Dendrogram grouping cell types according to gene expression pattern similarity. **C)** Schematic illustrating the concept of intercellular communication. Number of interactions indicates the variety of signaling pathways (e.g., signaling A [pink], signaling B [blue], signaling n) used between cell types A and B. Interaction strength implements the relative activity with which a specific signaling pathways (e.g., signaling A [pink]) is utilized between these cell types. Arrow thickness correlates with interaction strength. **D)** Number of interactions and interaction strength for healthy-1 (dark grey) and USH1C-1 (dark orange) ROs. **E)** Number and strength of interactions for all 18 cell types in healthy-1 and USH1C-1 ROs. Outgoing interaction strength is shown on x-axis. Incoming interaction strength is shown on y-axis. The number of distinct interactions is indicated by dot size.

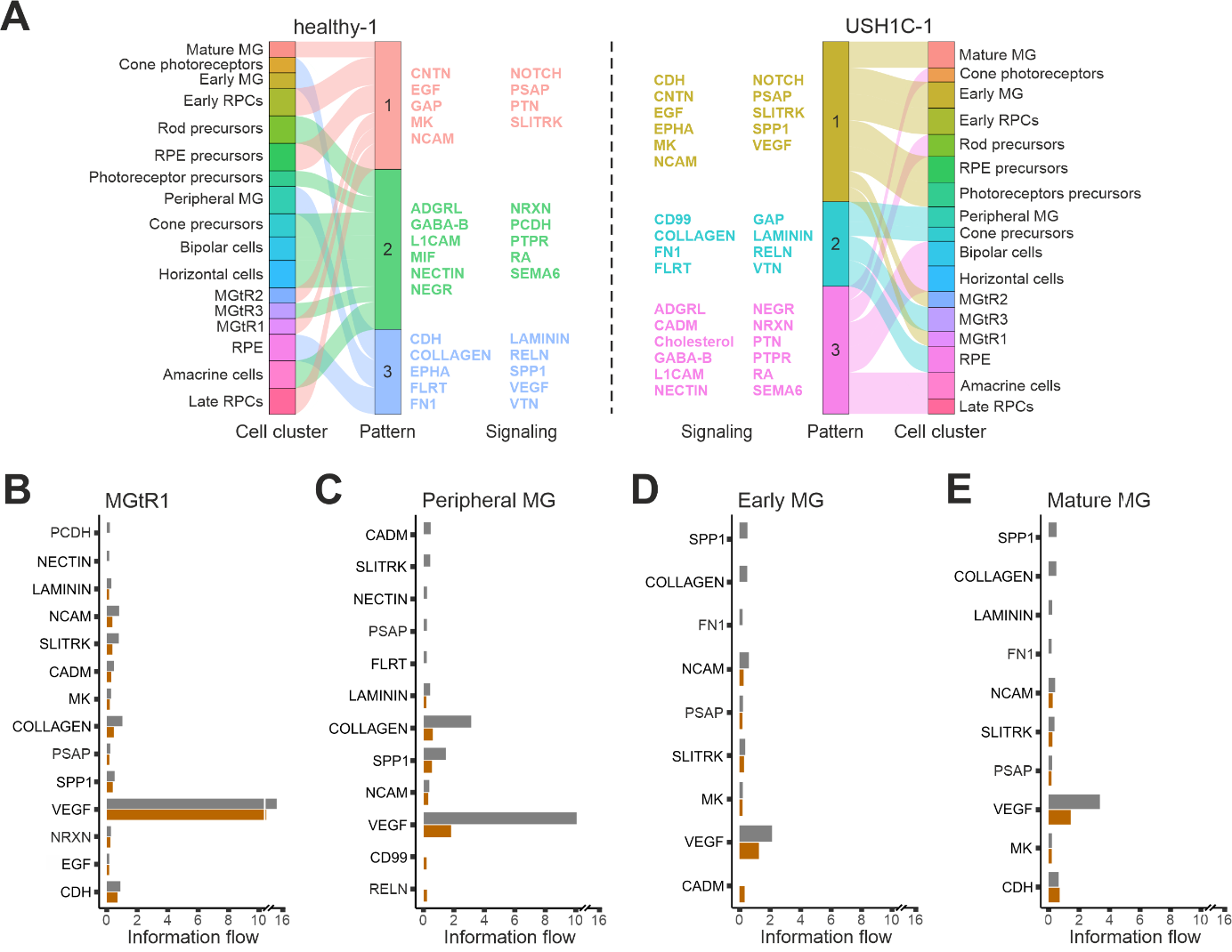

**Fig. S4 Incoming intercellular signaling in mature healthy-1 and USH1C-1 retinal organoids. A)** River plot grouping cell types of healthy-1 and USH1C-1 ROs according to similarities in their incoming communication. Each signaling pattern is defined by different signaling pathways. **B-E)** Comparison of significant differential incoming signaling pathways from MGtR1, peripheral MG, early MG and mature MG cells of healthy-1 (dark grey) and USH1C-1 (dark orange) ROs. Statistical analysis was done using Wilcoxon test.

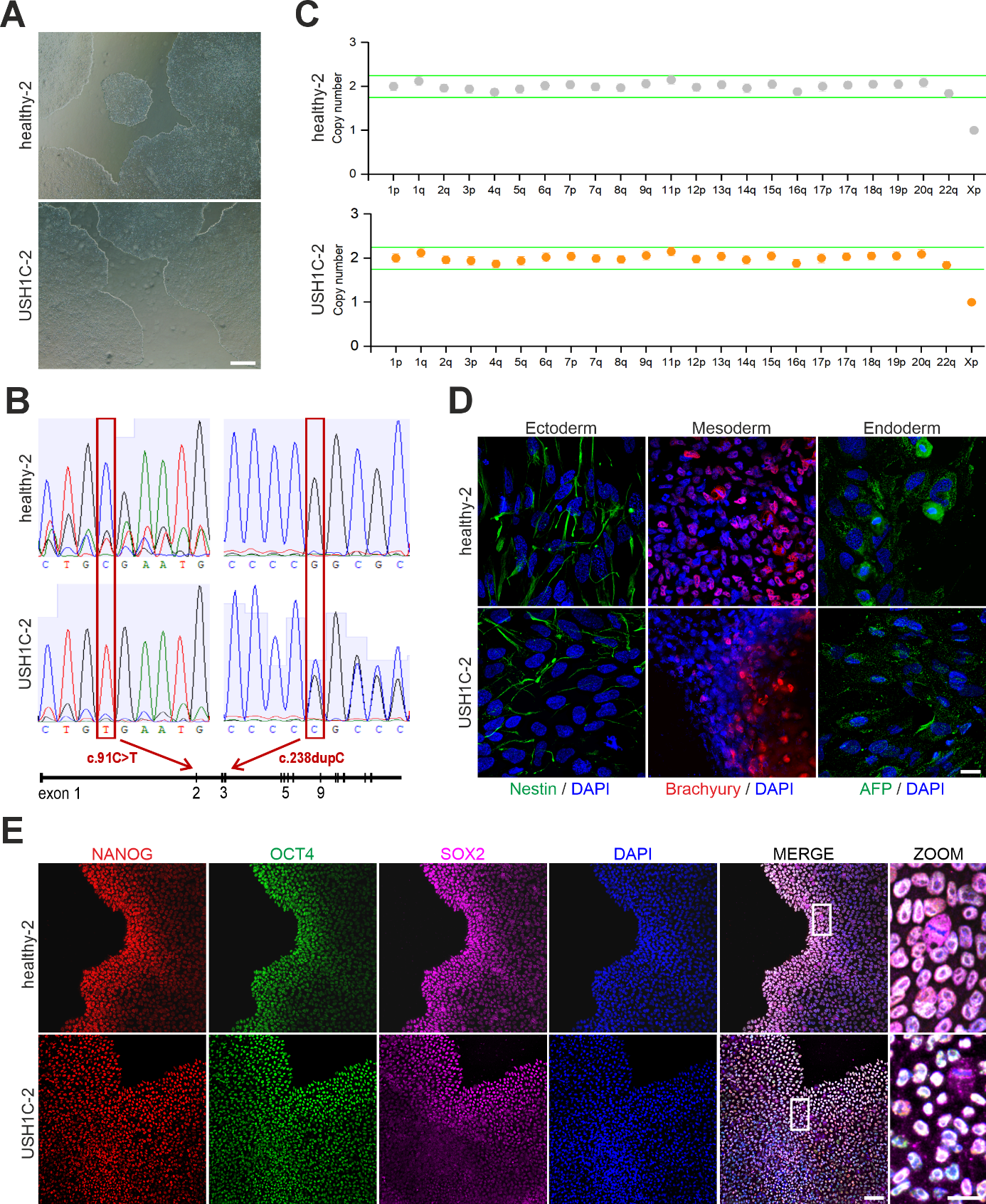

**Fig. S5 Characterization of healthy-2 and USH1C-2 human induced pluripotent stem cells. A)** Morphology of healthy-2 and USH1C-2 human induced pluripotent stem cell (hiPSC) colonies. Scale bar: 500 µm. **B)** Sanger sequencing determined the absence or presence of the c.91C>T; p.(R31*) and the c.238dupC; p.(R80Pfs*69) *USH1C* pathogenic variants in exon 2 and 3, respectively. **C)** Genomic stability of healthy-2 and USH1C-2 hiPSCs was assessed by testing for copy number variations (CNVs). **D)** Differentiation ability of healthy-2 and USH1C-2 hiPSCs into the three germ layers was assessed by an embryoid body (EB)-assay. Nestin was used as an ectodermal, Brachyury as a mesodermal and AFP as an endodermal marker. Scale bar: 25 µm. **E)** Expression of transcription factors NANOG, OCT4 and SOX2 confirmed the undifferentiated state of healthy-2 and USH1C-2 hiPSCs. Scale bars: 75 µm and 25 µm.

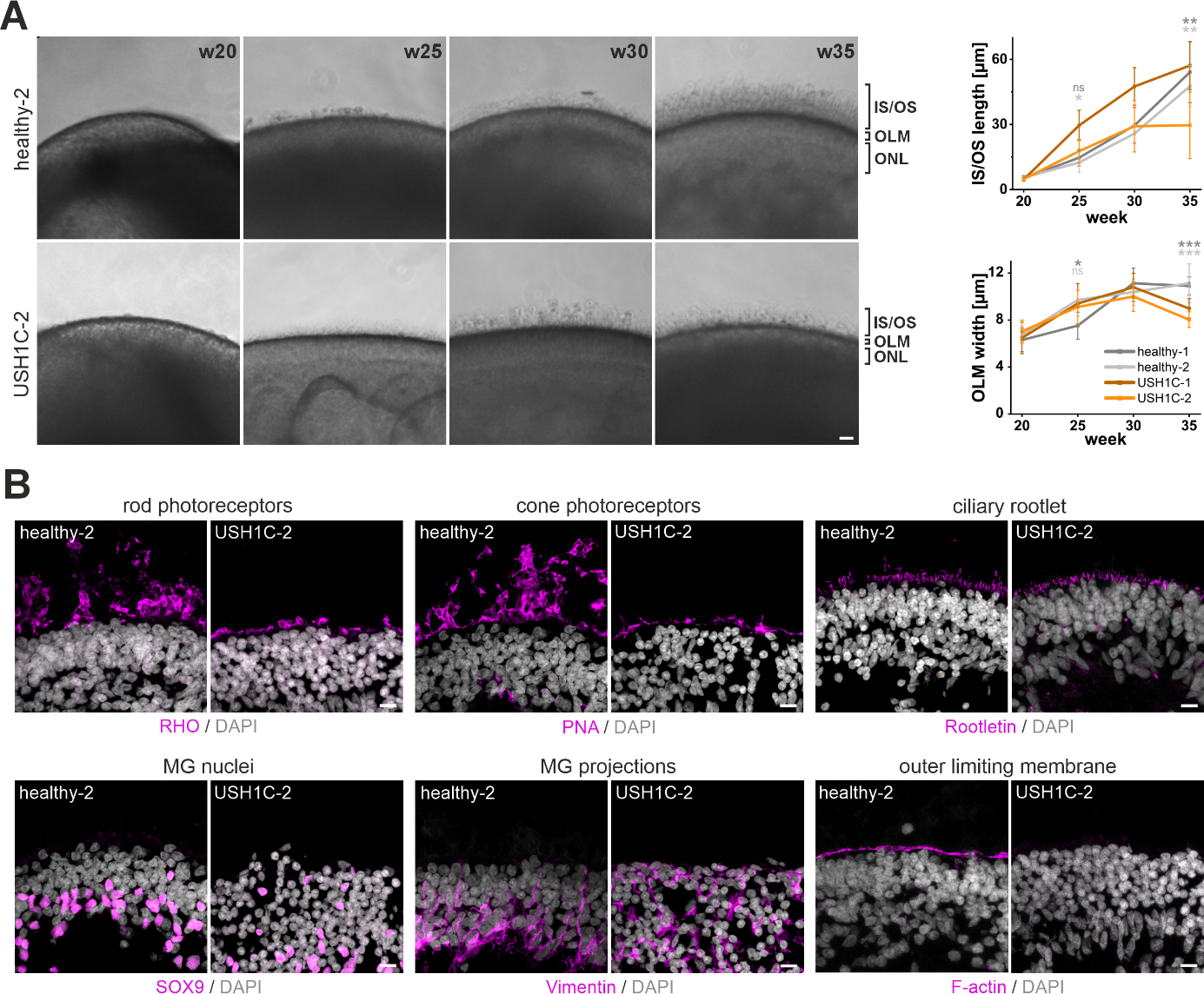

**Fig. S6 Morphological characterization of healthy-2 and USH1C-2 retinal organoids. A)** Representative images showing the final 15 weeks of healthy-2 and USH1C-2 RO maturation. Scale bar: 25 µm. Quantification of the IS/OS length and OLM thickness at w20, 25, 30 and 35. Independent samples *t*-test for n = 6-10 ROs per cell line/time point, 12 measurements per RO. Significant differences of USH1C-2 to healthy-1 and USH1C-2 to healthy-2 is indicated in dark grey and light grey asterix, respectively. **B)** IF-analysis of mature ROs for rod photoreceptors (RHO), cone photoreceptors (PNA), photoreceptor ciliary rootlet (Rootletin), MG cells (SOX9) and MG projections (Vimentin) and F-actin. Scale bar: 10 µm. n = 3 ROs.

**Table S1. Primers used for PCR.**

| **Primer name** | **Sequence 5'-3'** | **Amplicon [bp]** | **Experiment** |
| --- | --- | --- | --- |
| gDNA_harm_R31*_fwd | GGGTGGTCTGCATAGGTCTG | 858 | Sequencing of hiPSCs |
| gDNA_harm_R31*_rev | CCCTCGCAGTTGGAAAGGAA | 858 | Sequencing of hiPSCs |
| gDNA_harm_c238.dup_fwd | GGAAAGATCCCAACAGCCGA | 661 | Sequencing of hiPSCs |
| gDNA_harm_c238.dup_rev | CTGACCGCCTTTGATGAGGT | 661 | Sequencing of hiPSCs |

**Table S2. Primary antibodies used for immunofluorescence analysis.**

| **Primary Antibody** | **Protein** | **Host** | **Dilution** | **Supplier** | **Order Number** |
| --- | --- | --- | --- | --- | --- |
| Anti-AFP | Alpha fetoprotein | mouse | 1:200 | Sigma | WH0000174M1 |
| Anti-ARR3 | Arrestin 3 | goat | 1:200 | Novus Biologicals | NBP1-3700 |
| Anti-TBXT | Brachyury | goat | 1:200 | R&D Systems | AF2085 |
| Anti-CROCC | Rootletin | rabbit | 1:100 | selfmade | [1] |
| Anti-CTNNB1 | Catenin beta 1 | mouse | 1:100 | Santa Cruz | sc7963 |
| Anti-GNAT1 | G protein subunit alpha transducin 1 | rabbit | 1:250 | Proteintech | 55167-1-AP |
| Anti-GUCA1A | Guanylate cyclase activator 1A | rabbit | 1:200 | Proteintech | 12944-1-AP |
| Anti-NANOG | Nanog homeobox | goat | 1:200 | R&D Systems | AF1997 |
| Anti-NES | Nestin | mouse | 1:200 | Novus Biologicals | NBP1-92717SS |
| Anti-NRL | Neural retina leucine zipper | goat | 1:200 | R&D Systems | AF-2945-SP |
| Anti-OCT4A | POU class 5 homeobox 1 | rabbit | 1:400 | Cell Signaling | 2840 |
| Anti-RCVRN | Recoverin | rabbit | 1:300 | Millipore | AB5585 |
| Anti-RHO | Rhodopsin | mouse | 1:300 | Invitrogen | MA1-722 |
| Anti-SAG | S-antigen visual arrestin | mouse | 1:100 | selfmade | [2] |
| Anti-SOX2 | SRY-box transcription factor 2 | mouse | 1:500 | Santa Cruz | sc17320 |
| Anti-SOX9 | SRY-box transcription factor 9 | rabbit | 1:100 | Abcam | ab185966 |
| Anti-VIM | Vimentin | mouse | 1:200 | Abcam | sc32322 |

**Table S3. Secondary antibodies and dyes used for immunofluorescence analysis.**

| **Secondary Antibody and Dyes** | **Dilution** | **Supplier** | **Order Number** |
| --- | --- | --- | --- |
| ALEXA 488 donkey-anti-goat | 1:400 | Abcam | 150133 |
| CF 555 donkey-anti-goat | 1:400 | Sigma | SAB4600059 |
| ALEXA 488 donkey-anti-mouse | 1:400 | Invitrogen | A-21202 |
| ALEXA 647 donkey-anti-mouse | 1:400 | Invitrogen | A-31571 |
| ALEXA 488 goat-anti-rabbit | 1:400 | Invitrogen | A-11034 |
| CF 543 donkey-anti-rabbit | 1:400 | Biotrend | 20308-1 |
| CF 640R donkey-anti-rabbit | 1:400 | Biotrend | 20178 |
| 4’,6-diamidino-2-phenylindole (DAPI) | 1:12000 | Roth | 6335.1 |
| Peanut Agglutinin (PNA) | 1:400 | Invitrogen | L21409 |
| Rhodamin-Phalloidin | 1:400 | Sigma | P1951 |

**Table S4. Marker genes for UMAP representing 31 different cell types.**

See Excel file.

**Table S5. Marker genes for UMAP** **representing 18 different cell types.**

See Excel file.

**Table S6. Differential outgoing and incoming interaction strength per cell type.** A decreased cellular interaction in USH1C-1 ROs compared to healthy-1 ROs is highlighted in blue. An increased cellular interaction in USH1C-1 ROs compared to healthy-1 ROs is highlighted in red.

| **Cell type** | **Differential outgoing interaction strength** |  | **Differential incoming interaction strength** | **Cell type** |
| --- | --- | --- | --- | --- |
| MGtR1 | -1.43 |  | -1.55 | MGtR1 |
| Early MG | -0.91 |  | -1.49 | Peripheral MG |
| RPE precursors | -0.91 |  | -0.69 | Mature MG |
| Horizontal cells | -0.66 |  | -0.65 | Early MG |
| Peripheral MG | -0.48 |  | -0.48 | RPE precursors |
| MGtR2 | -0.43 |  | -0.30 | Horizontal cells |
| Amacrine cells | -0.39 |  | -0.29 | Amacrine cells |
| Mature MG | -0.36 |  | -0.26 | RPE |
| RPE | -0.34 |  | -0.25 | MGtR2 |
| Bipolar cells | -0.25 |  | -0.24 | Bipolar cells |
| MGtR3 | -0.24 |  | -0.22 | Late RPCs |
| Early RPCs | -0.17 |  | -0.17 | Rod precursors |
| Rod precursors | -0.11 |  | -0.13 | Rod photoreceptors |
| Rod photoreceptors | -0.11 |  | -0.12 | MGtR3 |
| Photoreceptor precursors | -0.05 |  | -0.09 | Early RPCs |
| Late RPCs | -0.03 |  | 0.01 | Cone precursors |
| Cone photoreceptors | 0.04 |  | 0.04 | Cone photoreceptors |
| Cone precursors | 0.05 |  | 0.10 | Photoreceptor precursors |

**Table S7. Differential number of interactions per cell type.** A decreased cellular interaction in USH1C-1 ROs compared to healthy-1 ROs is highlighted in blue. An increased cellular interaction in USH1C-1 ROs compared to healthy-1 ROs is highlighted in red.

| **Cell type** | **Differential number of interactions** |
| --- | --- |
| MGtR1 | -107 |
| MGtR2 | -52 |
| Mature MG | -51 |
| RPE precursors | -46 |
| Amacrine cells | -44 |
| Early MG | -39 |
| Early RPCs | -36 |
| Horizontal cells | -35 |
| Peripheral MG | -29 |
| Bipolar cells | -13 |
| RPE | -12 |
| MGtR3 | 3 |
| Rod photoreceptors | 6 |
| Photoreceptor precursors | 20 |
| Cone precursors | 23 |
| Late RPCs | 27 |
| Cone photoreceptors | 34 |
| Rod precursors | 34 |

**Table S8. Signaling pathways of MG subtypes related to retinal adhesion.** Signaling is associated with different protein families that fullfil functions related to ECM-receptor interaction, focal adhesion, cell adhesion and adherens junctions.

| **Signaling** | **Function** | **Literature** |
| --- | --- | --- |
| CADM | Cell adhesion molecules. | KEGG pathway database |
| CDH | Cell adhesion molecules, adherens junction. Tissue morphogenesis, neural circuit formation, neuronal survival, photoreceptor development and maintenance. CDH mutations are associated with cone-rod dystrophy, rod-cone dystrophy. CDH23 mutations are associated with Usher syndrome type 1D. | KEGG pathway database; [3] |
| CD99 | Cell adhesion molecules. | KEGG pathway database |
| CNTN | Cell adhesion molecules. Neural cell migration, axon guidance. | KEGG pathway database; [4] |
| COLLAGEN | ECM-receptor interaction. ECM component. Maintenance of structural strength of connective tissues. | KEGG pathway database; [5] |
| FLRT | Cell adhesion molecules. | [6] |
| FN1 | ECM-receptor interaction, regulation of actin cytoskeleton, Focal adhesion. Serves as a template for assembly of other ECM proteins. | KEGG pathway database; [7, 8] |
| LAMININ | ECM-receptor interaction. ECM Component. Maintenance of structural integrity. | KEGG pathway database; [8, 9] |
| NCAM | Cell adhesion molecules. Cell interaction, cell migration, axon growth, synaptic plasticity, regeneration. | KEGG pathway database; [10, 11] |
| NECTIN | Cell adhesion molecules, adherens junction. | KEGG pathway database |
| NRXN | Cell adhesion molecules. Synapse formation, synapse regulation. | KEGG pathway database; [12] |
| PCDH | Cell adhesion molecules. PCDH15 mutations are associated with Usher syndrome type 1D. | [13] |
| PTPR | Cell adhesion molecules, adherens junction. | KEGG pathway database |
| RELN | ECM-receptor interaction, focal adhesion. | KEGG pathway database; [14] |
| SLITRK | Cell adhesion molecules. Synapse formation. | KEGG pathway database; [15] |
| SPP1 | ECM-receptor interaction, focal adhesion. | KEGG pathway database |

**Table S9. DEG analysis for UMAP representing 18 different cell types.**

See Excel file.

**Table S10. Number of identified DEGs per cell type.**

| **Cell type** | **Number of DEGs** |
| --- | --- |
| Rod photoreceptors | 253 |
| Cone photoreceptors | 233 |
| Mature MG | 192 |
| RPE precursors | 110 |
| Bipolar cells | 98 |
| Early RPCs | 96 |
| Early MG | 81 |
| Rod precursors | 24 |
| Cone precursors | 23 |
| Horizontal cells | 19 |
| RPE | 12 |
| MGtR2 | 9 |
| Amacrine cells | 7 |
| Photoreceptor precursors | 7 |
| MGtR3 | 4 |
| Peripheral MG | 2 |
| Late RPCs | 1 |
| MGtR1 | 1 |

**Table S11. GO-term and KEGG analysis based on DEGs.**

See Excel file.

**Table S12. DEGs related to phototransduction in rod and cone photoreceptors.** DEGs associated with phototransduction in rod and cone photoreceptors were identified either automatically using ClueGO or manually via literature research. A drecreased gene expression in USH1C-1 ROs compared to healthy-1 ROs is highlighted in blue. An increased gene expression in USH1C-1 ROs compared to healthy-1 ROs is highlighted in red.

|  | **Rod photoreceptors** | **Cone photoreceptors** |
| --- | --- | --- |
| **Automatically identified DEGs related to phototransduction** | GNAT1 | GNAT2 |
|  | GRK1 | GUCA1B |
|  | PDE6A | PDE6B |
|  | PDE6G |  |
|  | RHO |  |
|  | SAG |  |
|  | GNGT1 |  |
|  | RCVRN |  |
| **Manually identified DEGs related to phototransduction** | GUCA1A | ARR3 |
|  | GNB3 | GNB3 |
|  | PDE6H | GNGT2 |
|  |  | OPN1MW3 |
|  |  | PDE6H |
|  |  | OPN1MW |
